# Distributed brain and spinal cord dynamics during micturition revealed by functional ultrasound imaging

**DOI:** 10.64898/2026.10.26.762042

**Authors:** Helma F. Zareian, Sofia Sakellaridi, Yu Tung Lo, Kevin Wu, Hui Zhong, Kofi Agyeman, Wooseong Choi, Aidin Abedi, Victor Reggie Edgerton, Evgeniy Kreydin, Darrin J. Lee, Charles Y Liu, Vassilios Christopoulos

**Affiliations:** Department of Biomedical Engineering, Viterbi School of Engineering, University of Southern California, Los Angeles, CA, USA; Casa Colina Hospital and Centers for Healthcare, Pomona, CA 91767, USA; Department of Neurological Surgery, Keck School of Medicine, University of Southern California, Los Angeles, CA, USA; Neurorestoration Center, Keck School of Medicine, University of Southern California, Los Angeles, CA, USA; Department of Neurosurgery, National Neuroscience Institute, Singapore; Rancho Los Amigos National Rehabilitation Center, Downey, CA, USA; Institute of Urology, Keck School of Medicine, University of Southern California, Los Angeles, CA, United States

## Abstract

Micturition relies on coordinated activity across distributed neural circuits spanning the brain and spinal cord, yet the spatiotemporal dynamics of these processes remain poorly understood. Disruption of these pathways in conditions such as spinal cord injury, Parkinson’s disease, multiple sclerosis, and aging can lead to neurogenic bladder dysfunction. Here, we used functional ultrasound imaging (fUSI), a modality with high spatiotemporal resolution and single-trial sensitivity, to investigate central nervous system activity during micturition. In anesthetized rats undergoing controlled bladder filling and volitional voiding, we recorded power Doppler signals from either the brain or the lumbosacral spinal cord. We observed distinct, region-specific hemodynamic responses associated with micturition across multiple brain regions, including the thalamus, hypothalamus, midbrain, and neocortex – particularly sensorimotor and retrosplenial areas – as well as dorsal and ventral spinal cord circuits. Furthermore, population-level activity in these regions enabled accurate decoding of bladder pressure dynamics. These results provide a systems-level characterization of central nervous system activity during micturition and establish fUSI as a platform for developing neural interfaces to restore bladder control.

## Introduction

Micturition is a complex process that involves pathways at many levels of the brain and spinal cord, ensuring appropriate storage and voiding of urine^1^. Diseases or injuries of the nervous system can cause the re-emergence of involuntary micturition, resulting in neurogenic bladder dysfunction, a significant contributor to morbidity, mortality, and reduction in quality of life. Understanding the functional circuits involved in micturition is crucial for addressing neurogenic bladder conditions and developing therapeutic strategies.

In the past few decades, rapid development of functional neuroimaging, including functional magnetic resonance imaging (fMRI)^2^, single-photon emission computed tomography (SPECT)^3^, and positron emission tomography (PET)^4,5^, has significantly contributed to our understanding of the neural mechanisms underlying micturition. However, many of these studies focus on the mechanisms involved in the urine storage phase. These studies involved full bladder filling or desire to induce voiding, by having subjects drink water^3,5,6^, perform repeated saline perfusion and withdrawal through urinary catheters^7,8^, or engage in repeated spontaneous contractions of pelvic floor muscles to activate relevant brain regions^9–11^ (for review see^12^). Conversely, few studies have explored the voiding phase, primarily due to two challenges: (1) the low sensitivity of traditional imaging methods that necessitates multiple trials (i.e., participants are required to urinate several times during the experiment) to achieve adequate signal-to-noise ratio, and (2) many participants report discomfort when urinating in a controlled experimental setting, leading to a higher rate of exclusions. As a result, our understanding of the neural basis of micturition – particularly the transition from storage to voiding – remains incomplete.

Beyond the brain, the spinal cord plays a critical role in coordinating bladder function, yet investigations into spinal cord activity during micturition are even more limited. This gap in research is largely due to anatomical and technical challenges^13–16^. In particular, the bony, fascial enclosure and small cross-sectional dimensions of the spinal cord combined with susceptibility artifacts due to local magnetic field inhomogeneities generated by interfaces between surrounding bones, ligaments, soft tissues and cerebrospinal fluid (CSF) make the spinal cord an unfavorable target for traditional neuroimaging techniques ^13–16^. These challenges point to a dire need for neuroimaging technologies capable of capturing brain and spinal cord activity during micturition with high spatiotemporal resolution and sensitivity.

Functional ultrasound imaging (fUSI) is an emerging modality with exceptional capabilities, offering high resolution (∼100 μm and ∼100 ms), extensive penetration depth (up to 8 cm), ability to measure slow blood flow (as low as 1 mm/s), and sensitivity to detect hemodynamic changes of only 2% within a single trial^17–20^. Based on power Doppler (pD) signal, fUSI measures changes in blood volume by detecting backscattered echoes from red blood cells moving within its field of view. While fUSI is a hemodynamic technique, its superior spatiotemporal performance and single-trial sensitivity offer a substantially closer connection to the underlying neuronal signals than achievable with other hemodynamic methods such as fMRI.

In the current study, we used fUSI to investigate the brain and spinal cord mechanisms underlying micturition. We recorded fUSI images from the brain and spinal cord in anesthetized rats during controlled bladder filling, which elicited spontaneous voiding. We identified region-specific hemodynamic changes across both brain and spinal cord structures during micturition, including void-related activation patterns and responses associated with bladder pressure fluctuations. Notably, the diencephalon, midbrain, and neocortex – particularly the sensorimotor and retrosplenial cortices – exhibited strong activation, along with the dorsal and ventral regions of the lumbosacral spinal cord. We further demonstrated that pD signals from both brain and spinal cord can accurately predict bladder pressure dynamics.

Together, these findings provide a systems-level characterization of central nervous system activity during micturition, extending prior human spinal cord fUSI studies of micturition^21^, and revealing distributed hemodynamic activity associated with the transition from bladder storage to voiding. More broadly, this work advances our understanding of the neural circuits underlying bladder control and provides a framework for investigating autonomic dysfunction and developing neurotechnology-based therapeutic strategies for neurogenic bladder disorders.

## Results

To characterize brain and spinal cord hemodynamic responses during micturition, we simultaneously acquired pD images and continuous bladder pressure recordings in anesthetized rats (**Figure 1A**). We collected pD data from the brain (n = 5) and spinal cord (n = 7) during urodynamically-controlled bladder filling cycles interspersed by a hold period and followed by a washout period (**Figure 1D**). Functional ultrasound recordings were acquired using a 128-channel linear array miniaturized ultrasound transducer (Iconeus One) with 15.6-MHz center frequency. The probe was mounted on a three-axis motorized positioning system and oriented in the sagittal plane. For brain imaging, craniectomies extended from bregma to foramen magnum, with lateral borders defined by the superior temporal lines (**Figure 1B**). For spinal cord imaging, the probe was aligned to span the laminectomy window between the T12 and L1 vertebrae, exposing spinal levels L2-S1 (**Figure 1C**).

**Figure 1.**
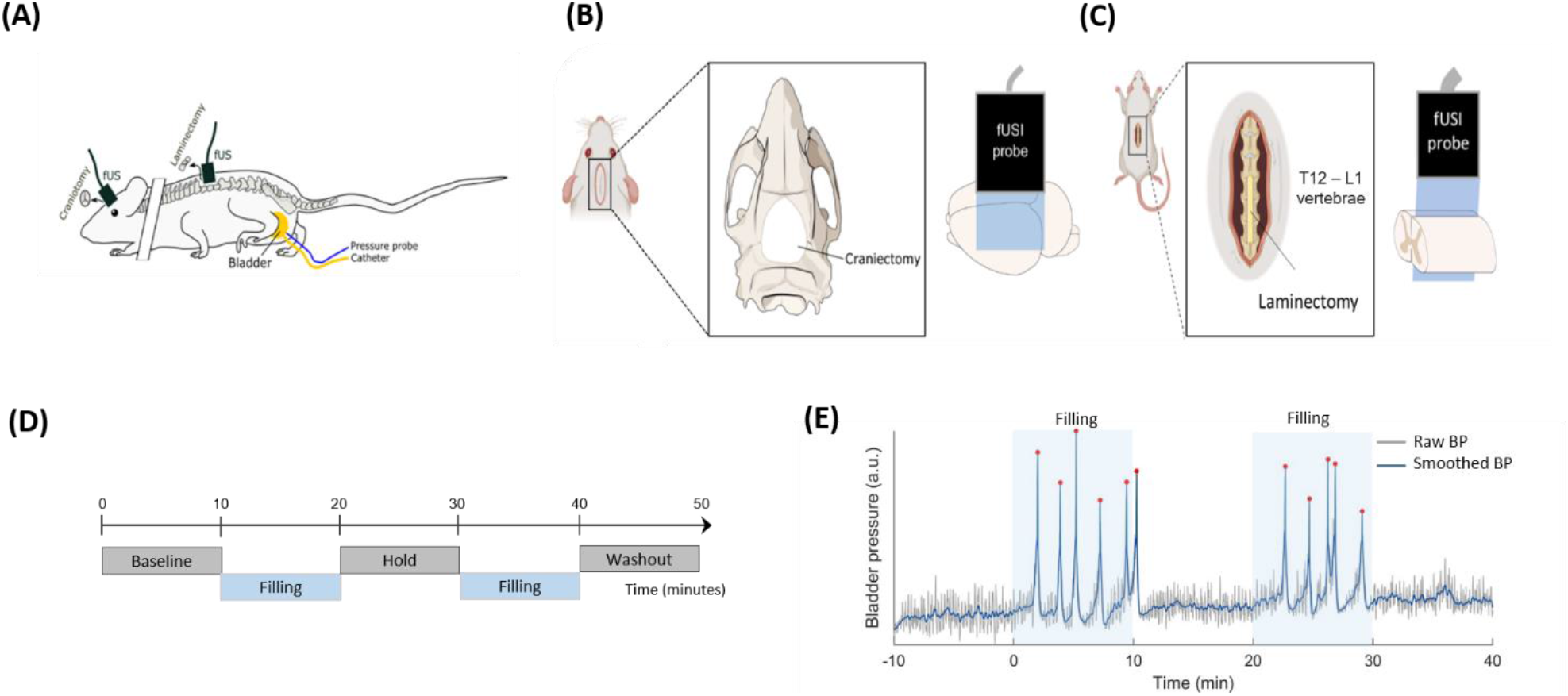
Experimental setup for simultaneous fUSI acquisition and urodynamic recordings. **(A)** Simultaneous urodynamic and fUSI recordings performed over the brain and spinal cord in experimental rats (brain and spinal cord fUSI recordings were acquired in separate animal groups). **(B)** Configuration of the craniotomy window and approximate placement coverage of the fUSI probe on the brain. With the craniectomy window spanning from bregma to rostral to the foramen magnum and lateral borders at the superior temporal line. **(C)** Similar to panel B, but for the laminectomy in the spinal cord, spanning from the T12-L1 vertebrae (**D**) The urodynamic protocol of bladder filling and hold, consisting of 10 min baseline activity recording, followed by sequential cycles of bladder filling with 0.9% saline at a constant rate of 0.1 ml/min, interleaved with hold periods, and concluded with 10 min of washout recording. **(E)** Representative bladder pressure traces recorded during the 50 min experimental session. *Gray curve*: raw bladder pressure signal, including two 10 min filling periods during continuous saline infusion (0.1 ml/min). Red dots indicate bladder pressure peaks identified using threshold-based peak detection. *Blue curve*: bladder pressure trace after Gaussian smoothing to reduce high-frequency noise while preserving the overall signal profile and the local structure surrounding voiding peaks.

Each experiment lasted 50 minutes and consisted of 10 minutes of baseline recordings followed by two filling phases separated by an intermediate hold period and ending with a 10-minute washout period, designed to induce spontaneous voiding (**Figure 1D**). Continuous intravesical pressure was recorded using a pressure transducer and urinary catheter simultaneously with fUSI of the brain or spinal cord. **Figure 1E** shows a representative bladder pressure trace during the filling and holding phases, highlighting the sharp acceleration and deceleration patterns characteristic of voiding events (red dots).

The craniectomy configuration provided us access to key brain regions, including the sensorimotor cortex, retrosplenial cortex, hippocampal formation, diencephalon, midbrain, rostral parts of the cerebellum, and brainstem (**Figure 2A**). Following laminectomy, major spinal cord vessels – including the anterior spinal artery (ASA), posterior spinal vein (PSV), and central sulcal artery (CSA) – were clearly identifiable within the imaging field of view (**Figure 2B**).

**Figure 2.**
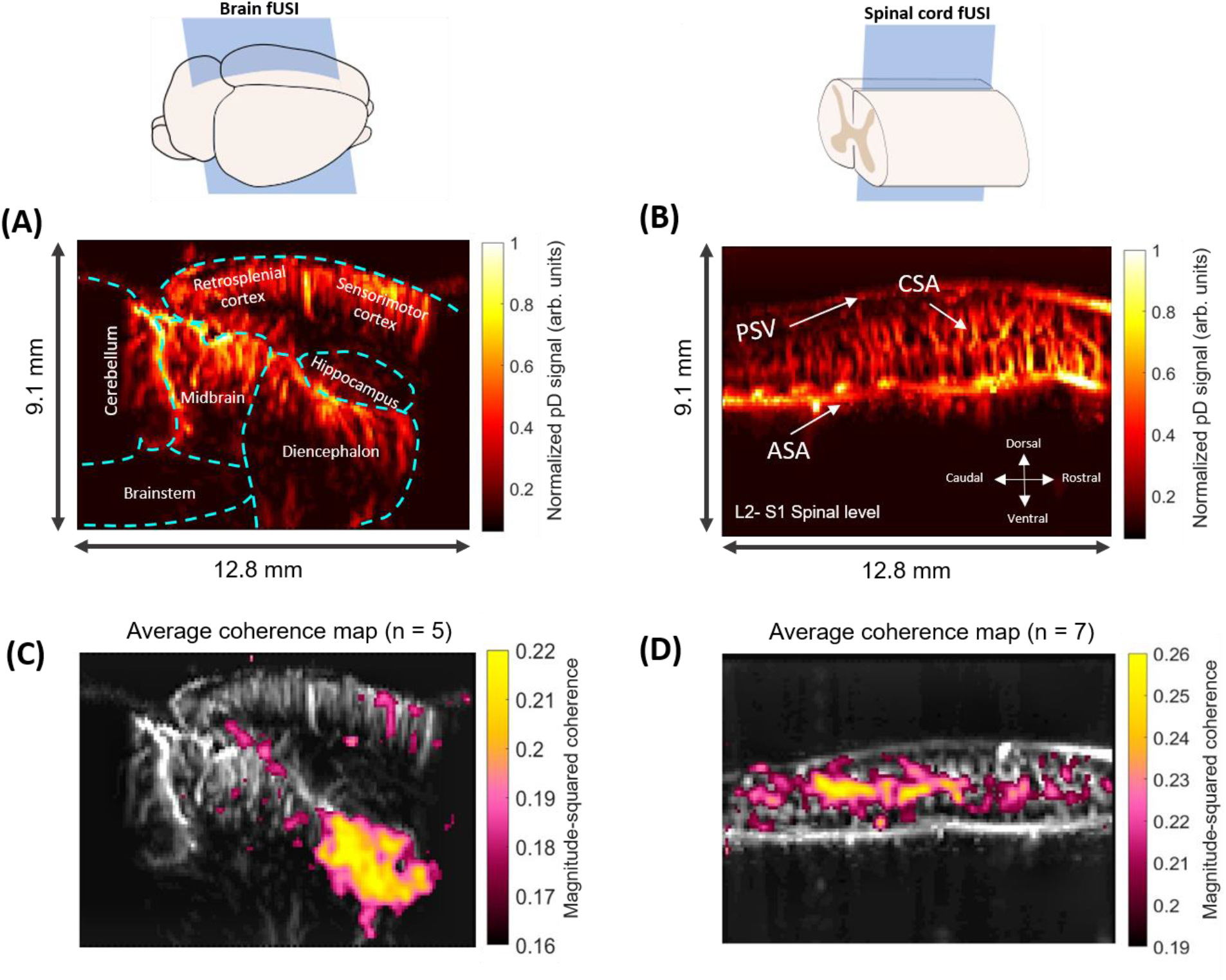
Pixel-wise coherence analysis between bladder pressure and power Doppler (pD) signals. **(A)** Representative pD vascular map of a rat brain illustrating vascular anatomy and major anatomical regions. **(B)** Representative pD vascular map of the rat spinal cord. ASA: anterior spinal artery; PSV: posterior spinal vein; CSA: central sulcal artery**. (C)** Group-average coherence map for brain fUSI recordings (n = 5), thresholded to display regions within the top 10% of coherence values. **(D)** Group-average coherence map for spinal cord fUSI recording (n = 7), showing regions within the top 10% of coherence values. Coherence maps illustrate the spatial distribution of magnitude-squared coherence between bladder pressure fluctuations and pD signal dynamics within the 0.0033-0.033 Hz frequency range.

### Pixel-wise coherence analysis

To identify brain and spinal cord regions with hemodynamic activity that co-varied with bladder pressure fluctuations, we performed pixel-wise magnitude-squared coherence (MSC) analysis between pD signals and bladder pressure, focusing on the 0.0033-0.033 Hz frequency range (corresponding to periods of 30-300 s that span typical voiding cycles). This analysis quantifies the similarity of frequency profiles between hemodynamic and bladder-pressure signals, independent of phase, providing a frequency-domain map of their coupling across the entire recording period.

To generate group-level pixel-wise coherence maps, we performed inter-animal pD image registration to align individual vascular maps to common anatomical references. Brain pD images from rats 1–5 were registered to the reference animal (rat 1) using an affine transformation, whereas spinal cord pD images from rats 6–12 were aligned to rat 8 using a nonlinear locally weighted mean (LWM) transformation. This approach enabled the construction of average coherence maps across animals for both brain and spinal cord datasets.

In the brain recordings (n = 5), the group-averaged coherence map, thresholded to display the top 10% of coherence values, revealed a prominent cluster of strong coupling between bladder pressure and hemodynamic signal in the diencephalon, including the thalamic and hypothalamic areas (**Figure 2C**). Additional clusters were observed in sensorimotor cortical areas and midbrain structures, suggesting coordinated coupling between regional hemodynamic activity and intravesical pressure dynamics.

In the spinal cord recordings (n = 7), coherence analysis revealed strong coupling along the rostrocaudal axis from L2 to S1, with peak values localized primarily in the caudal-dorsal regions of the lumbosacral segments (**Figure 2D**). This coherence map likely reflects frequency-dependent coupling between lumbosacral circuits and bladder pressure fluctuations. Together, these findings indicate that both brain and spinal cord activity are temporally coupled to intravesical pressure dynamics during the micturition cycle.

### Spatiotemporal hemodynamic changes during micturition

To better characterize hemodynamic changes in the brain and spinal cord during the transition from bladder storage to voiding, we performed dynamic statistical parametric map (dSPM) analysis. Percent changes in blood volume (i.e., pD signal) in the brain (ΔCBV%) and spinal cord (ΔSCBV%) were computed using a non-overlapping 3-s time sliding window spanning ±16 s around the bladder pressure peak (**Figure 3A, 4A**). Activation maps revealed spatially distributed regions exhibiting significant increases (positive responses, reddish areas) and decreases (negative responses, bluish areas) in the pD signal across the peri-void period (**Figure 3B, 4B**). Each displayed dSPM frame in **Figure 3B** and **4B** represents the mean activation averaged across a 3-s time window, yielding 11 discrete steps spanning the full cycle. These responses were time-locked to the bladder pressure peak and evolved dynamically before, during, and after voiding.

**Figure 3.**
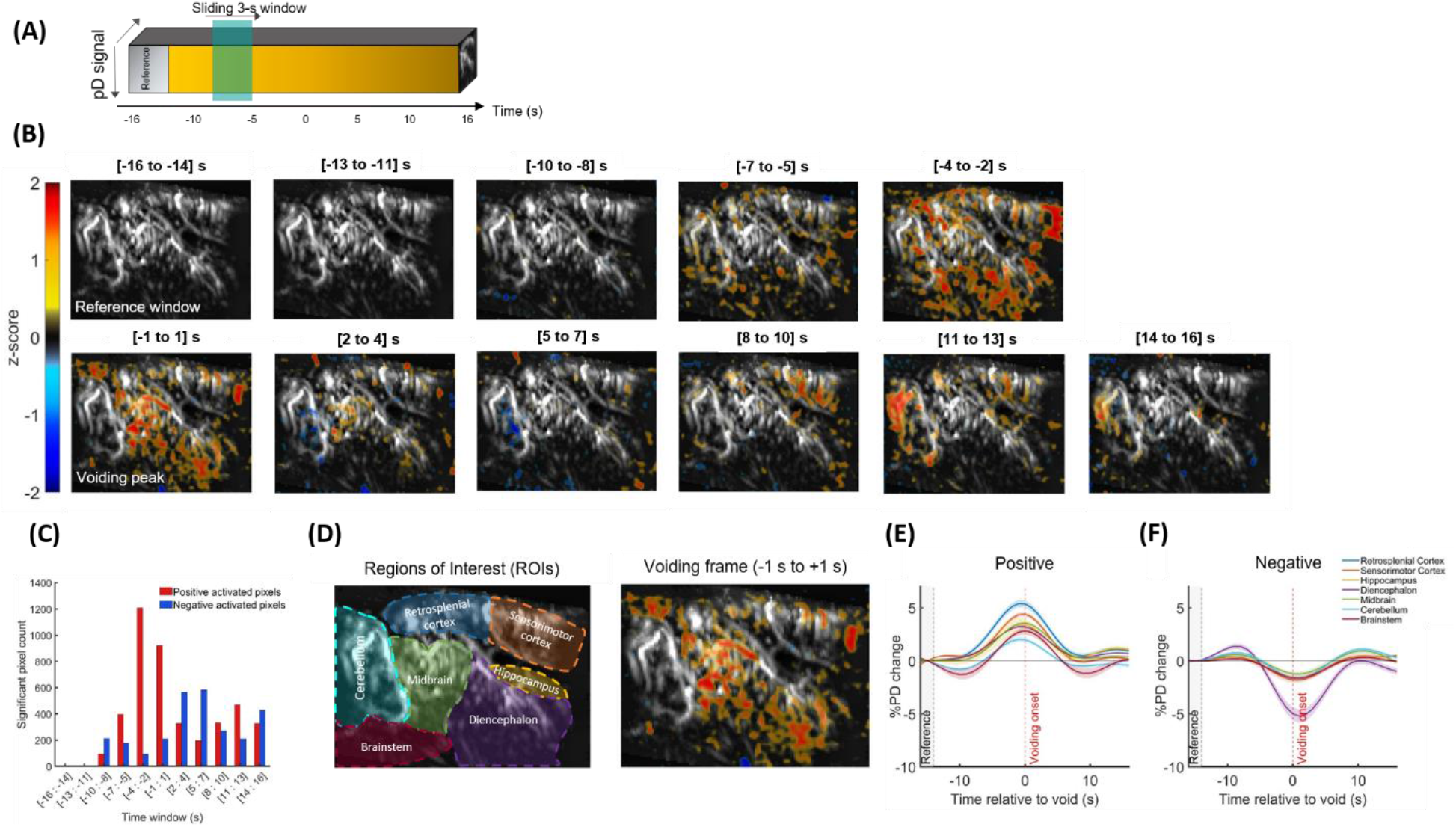
Brain dynamic statistical parametric maps (dSPMs) and ROI-specific event-related average (ERA) waveforms of the cerebral blood volume changes (ΔCBV) around the voiding peak. (A) dSPMs were generated using two-sided pixel-wise Student’s t-tests (p < 0.01), with false discovery rate (FDR) correction across image pixels, applied within non-overlapping 3-s sliding windows relative to a 3-s baseline reference period (−16 to −14 s). (B) Group-average activation maps (n = 5) showing brain regions exhibiting significant positive (red) or negative (blue) ΔCBV changes throughout the peri-void interval (−16 to +16 s relative to the voiding peak). For each pixel and time window, z-scores were calculated from percentage signal changes relative to the reference period. Statistical maps were overlaid onto the mean baseline pD image. For visualization purposes, all significantly modulated pixels were retained, low-amplitude z-scores were rendered transparent to reduce visual crowding, and Gaussian smoothing (σ = 1) was applied. **(C)** Quantification of the temporal evolution of voiding-related hemodynamic responses, measured as the number of pixels within the top 15% of significantly increased (red) and decreased (blue) ΔCBV values across the voiding cycle. (D) *Left panel*: Anatomically defined regions of interest (ROIs), including the retrosplenial cortex, sensorimotor cortex, hippocampal formation, diencephalon, midbrain, cerebellum and brainstem. *Right panel*: ROIs were selected from the dSPM frame centered on the voiding peak (−1 to +1 s relative to the voiding peak). **(E)** ERA waveforms corresponding to positively modulated pixels within the peak-centered dSPM frame (D, right), demonstrating robust ΔCBV increases surrounding the voiding peak. ERA waveforms were calculated as the mean activity across the top 15% of activated pixels within each ROI. **(F)** Same as panel E, but for the negatively modulated pixels. Retrosplenial cortex and hippocampal formation are not shown because these regions contained fewer than five negatively modulated pixels, precluding reliable estimation of negative ERA waveforms.

**Figure 4.**
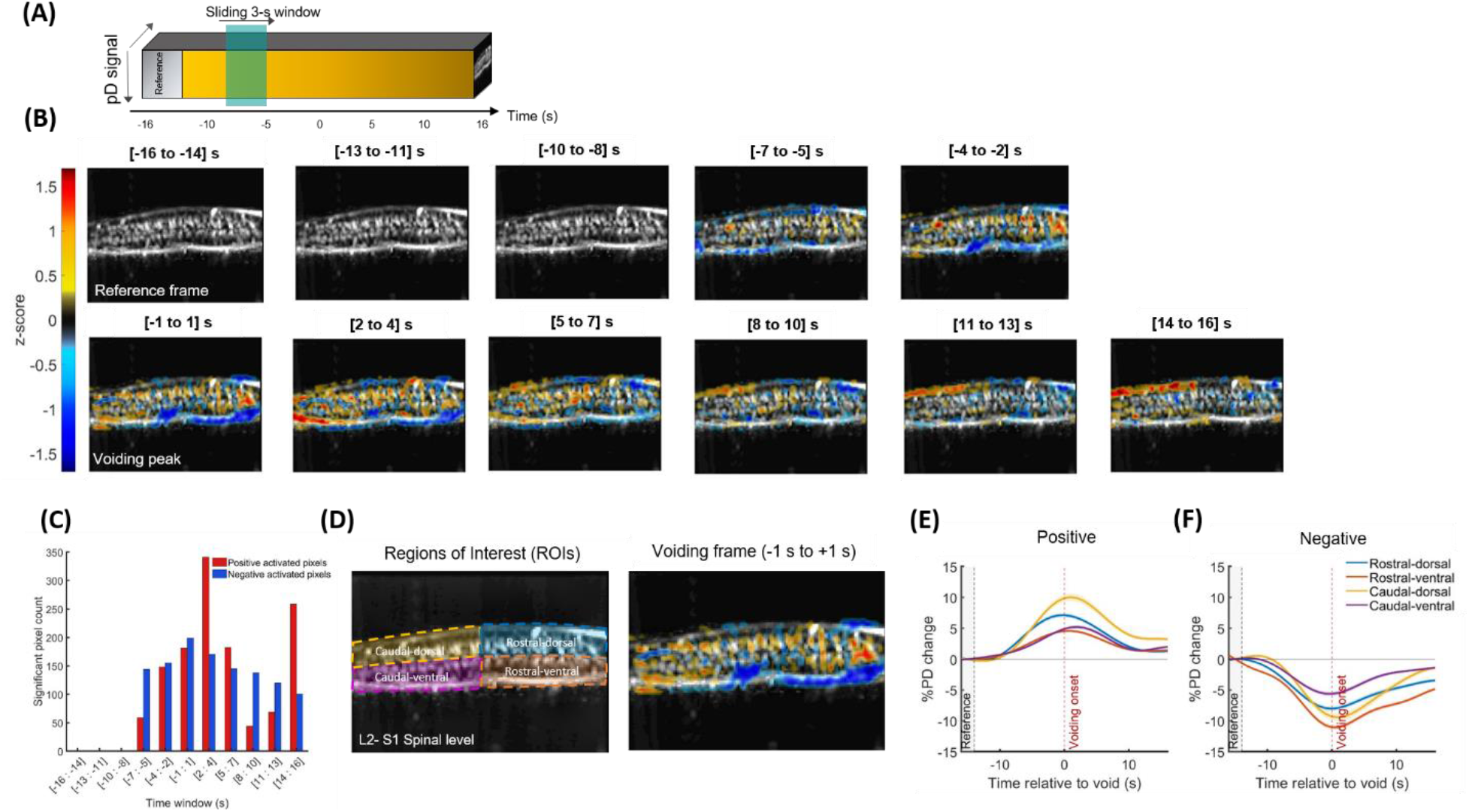
Spinal cord dynamic statistical parametric maps (dSPMs) and ROI-specific event-related average (ERA) waveforms of the spinal cord blood volume changes (ΔSCBV) around the voiding peak. **(A)** dSPMs were generated using two-sided pixel-wise Student’s t-tests (p < 0.01), with false discovery rate (FDR) correction across image pixels, applied within non-overlapping 3-s sliding windows relative to a 3-s baseline reference period (−16 to −14 s). **(B)** Group-average activation maps (n = 7) showing spinal cord regions exhibiting significant positive (red) or negative (blue) ΔSCBV modulation throughout the peri-void interval (−16 to +16 s relative to the voiding peak). For each pixel and time window, z-scores were computed from percentage signal changes relative to the reference period. Statistical maps were overlaid onto the mean baseline pD image of the reference rat (Rat 8). For visualization purposes, all significantly modulated pixels were retained, low-amplitude z-scores were rendered transparent to reduce visual crowding, and Gaussian smoothing (σ = 1) was applied. **(C)** Quantification of the temporal evolution of voiding-related spinal hemodynamic responses, measured as the number of pixels within the top 15% of significantly increased (red) and decreased (blue) ΔSCBV values across the voiding cycle. **(D)** *Left panel*: Anatomically oriented regions of interest (ROIs) in the spinal cord. The field of view (L2–S1 spinal levels) was divided along dorsal-ventral and rostral-caudal axes into four non-overlapping ROIs. *Right panel*: ROIs were selected from the dSPM frame centered on the voiding peak (−1 to +1 s relative to the voiding peak). **(E)** ERA waveforms corresponding to positively modulated pixels within the peak-centered dSPM frame (D, right), demonstrating robust ΔSCBV increases surrounding the voiding peak. ERA waveforms were calculated as the mean activity across the top 15% of activated pixels within each ROI. **(F)** Same as panel E, but for the negative modulated pixels.

### Dynamic brain activation pattern during micturition

**Figure 3B and Supplementary Movie 1** illustrate the spatiotemporal evolution of brain hemodynamics during the micturition cycle – i.e., transition from storage to voiding. No significant activation was detected during the reference and early pre-void period (−16 to −11 s relative to the voiding peak). As the voiding peak approached (-10 to -2 s), multiple brain regions exhibited significantly increased activity, including the retrosplenial and sensorimotor cortices, midbrain, diencephalon, hippocampus, and cerebellum. Near the voiding peak (-1 to +1 s), activation became more spatially localized, predominantly within the midbrain, diencephalon, and sensorimotor cortex. Following the peak (+2 to +7 s), activity decreased across most regions, accompanied by a significant reduction in pD signal in the cerebellum and a decline in the diencephalon and brainstem.

A secondary increase in activity was observed during the post-void period (+8 to +13 s) particularly within the sensorimotor cortex and cerebellum, indicating sustained hemodynamic modulation beyond the immediate voiding phase. At the late post-void period (+14 to +16 s), a transient decrease in activity was observed in the midbrain and diencephalon. Together, these temporal dynamics suggest a sequential recruitment and recovery of cortical, subcortical, and cerebellar networks involved in bladder sensation, coordination, and post-void recovery.

To further characterize the temporal evolution of voiding-related brain activation, we quantified the number of significantly activated and deactivated pixels (z < −1.35 or z > +1.582) at each dSPM frame across the full peri-void period (−16 to +16 s relative to voiding peak; **Figure 3C**). This analysis provides a temporal measure of the spatial extent of activation, complementing the visual patterns observed in the maps. The results showed that the number of positively activated pixels increased progressively prior to voiding, peaking at -4 to -2 s relative to the pressure maximum (i.e., voiding), followed by a decline and a secondary increase during the late post-void period. In contrast, negatively modulated pixels were less prevalent and exhibited a delayed rise, emerging primarily after the voiding peak (+2 to +7 s).

### Region-specific brain hemodynamic patterns associated with micturition

Next, we examined region-specific hemodynamic changes during micturition. To this end, we computed event-related averages (ERAs) of the brain pD signal changes (ΔCBV) for anatomically defined regions of interest (ROIs; **Figure 3D**). ROIs were outlined approximately according to anatomical boundaries defined in the Paxinos and Watson Rat Brain Atlas^22^ and included the retrosplenial cortex, sensorimotor cortex, hippocampal formation, diencephalon, midbrain, rostral parts of the cerebellum, and brainstem (**Figure 3D**, Left panel). For each ROI, ERA curves represent the mean percent change in the pD signal for the top 15% of significantly modulated positive and negative pixels in the frame centered on the voiding peak (**Figure 3D**, Right panel) relative to the 3-s reference window (−16 to −14 s). These ROI-specific ERAs reveal distinct temporal profiles of cerebral hemodynamic responses during the micturition cycle (**Figure 3E, 3F)**.

To compare ERA peak amplitudes across ROIs, we performed pairwise nonparametric Wilcoxon rank-sum tests, followed by Benjamini–Hochberg correction for multiple comparisons. Among the examined regions, the retrosplenial and sensorimotor cortices exhibited stronger pre-void positive responses than other ROIs (**Figure 3E**). Both cortices showed significantly greater responses compared with the diencephalon, midbrain, cerebellum and brainstem (retrosplenial: *p* ≤ 1.214 ×10^-6^ ; sensorimotor: *p* ≤ 0.0012). We next examined negative ERA responses across ROIs (**Figure 3F**). The retrosplenial cortex and hippocampal formation were excluded from this analysis because they contained fewer than five significantly negative pixels, making negative ERA estimates in these regions unreliable. Among the other regions, the diencephalon exhibited the strongest negative response, which was significantly greater than those observed in the midbrain, cerebellum, and brainstem (Wilcoxon rank-sum tests, *p* ≤ 0.0047).

### Dynamic spinal cord activation pattern during micturition

A similar dSPM analysis was performed in the spinal cord (**Figure 4B; Supplementary Movie 2**), revealing coordinated spatiotemporal patterns of hemodynamic activity across the L2-S1 segments. No significant changes in pD signals were observed during the early pre-void dSPM period (−13 to −8 s). During the late pre-void phase (-7 to -2 s), positive activation emerged predominantly in rostral spinal segments, accompanied by concurrent decreases in ventral (anterior spinal artery) and dorsal (posterior spinal vein) compartment. At the voiding peak (-1 to +1 s), activation extended caudally and became more pronounced within the ventral regions, while the negatively modulated rostroventral cluster increased in magnitude. Immediately following voiding (+2 to +4 s), positive activity became more prominent in the caudal segments of the spinal cord, whereas rostral regions exhibited predominantly negative responses, indicating spatially distinct patterns of activation and deactivation along the rostrocaudal axis.

During post-void period (+5 to +13 s), positive activity in the caudal spinal cord gradually shifted from ventral to dorsal compartments and continued to strengthen over time, reaching higher magnitude during the late post-void period (+14 to +16 s). In contrast, negative responses in rostral regions gradually diminished over this period. Overall, this spatiotemporal progression reflects a rostral-to-caudal propagation of hemodynamic activity across dorsal and ventral laminae.

To further understand the temporal dynamics of voiding-related spinal cord activity, we quantified the number of significantly modulated pixels (z < −1.488 or z > +1.537) at each dSPM frame across the peri-void interval, providing a time-resolved measure of spinal activity throughout the micturition cycle. As shown in **Figure 4C**, positively activated pixels peaked at +2 to +4 s relative to voiding peak, followed by a smaller late increase predominantly in caudal-dorsal regions (+14 to +16 s). In contrast, negatively activated pixels increased earlier (-1 to +1 s), primarily within rostral-ventral segments, before gradually declining.

Together, these results demonstrate that spinal cord activity is spatially and temporally organized across the micturition cycle, with distinct regional patterns of activation or deactivation observed before, during, and after voiding, and with sustained engagement of caudal regions extending beyond voiding completion.

### Region-specific spinal cord hemodynamic patterns associated with micturition

We further investigated region-specific hemodynamic dynamics by quantifying spinal cord pD signal changes (ΔSCBV) within anatomically oriented ROIs (**Figure 4D**). The field of view was partitioned along the dorsoventral and rostrocaudal axes, yielding four non-overlapping ROIs. This segmentation enabled assessment of spatial differences in hemodynamic responses across spinal subregions during the micturition cycle. As shown in **Figure 4E** and **4F**, ERA waveforms revealed distinct patterns along both the rostrocaudal and dorsoventral axes within the L2-S1 segments. Specifically, caudal-dorsal segments exhibited the strongest positive response (**Figure 4E)**, which was significantly greater than that observed in the other three ROIs (*p* ≤ 0.0017). For negative ERA responses, both the caudal-dorsal and rostral-ventral regions showed pronounced decreases (**Figure 4F)**. The rostral-ventral region exhibited significantly larger negative responses compared to the other three ROIs (*p* ≤ 0.019). These deactivations in spinal cord regions may reflect inhibitory gating within local spinal circuits. Overall, this analysis highlights spatial heterogeneity in both the amplitude and temporal evolution of hemodynamic responses across both cerebral and spinal networks during micturition.

### Decoding bladder pressure from brain and spinal cord hemodynamic signals

Next, we assessed whether pD signal from the brain and spinal cord can predict bladder pressure dynamics, a key step toward developing brain and spinal cord machine interface technologies for treating bladder dysfunction. Briefly, each 2D pD image (*N* × *M*) at time point *t* was reshaped into a 1D vector by flattening the image matrix (**Figure 5**). These vectors were then concatenated over time to form a 2D (pixels × time) matrix, with each row corresponding to a time point and each column represents an individual pixel. A separate vector contained the corresponding bladder pressure (BP) signal. We then trained a random forest regression (RFR) model – i.e., a supervised machine learning technique that leverages an ensemble of decision trees – to predict BP from the pD data. This method is well suited for the present application due to its ability to capture non-linear relationships, reduce overfitting, and scale efficiently to high-dimensional data. To determine optimal model complexity, we performed a grid search across the number of trees (*NumLearningCycles*) and minimum leaf size (*MinLeafSize*) and evaluated prediction performance across animals. Averaged performance across animals identified optimal decoding hyperparameter settings for the brain and spinal cord groups. For the brain model, optimal performance was achieved with *100* trees and a minimum leaf size of *50*, whereas the spinal cord model required *500* trees and a minimum leaf size of *100 (see Methods section for more details)*.

**Figure 5.**
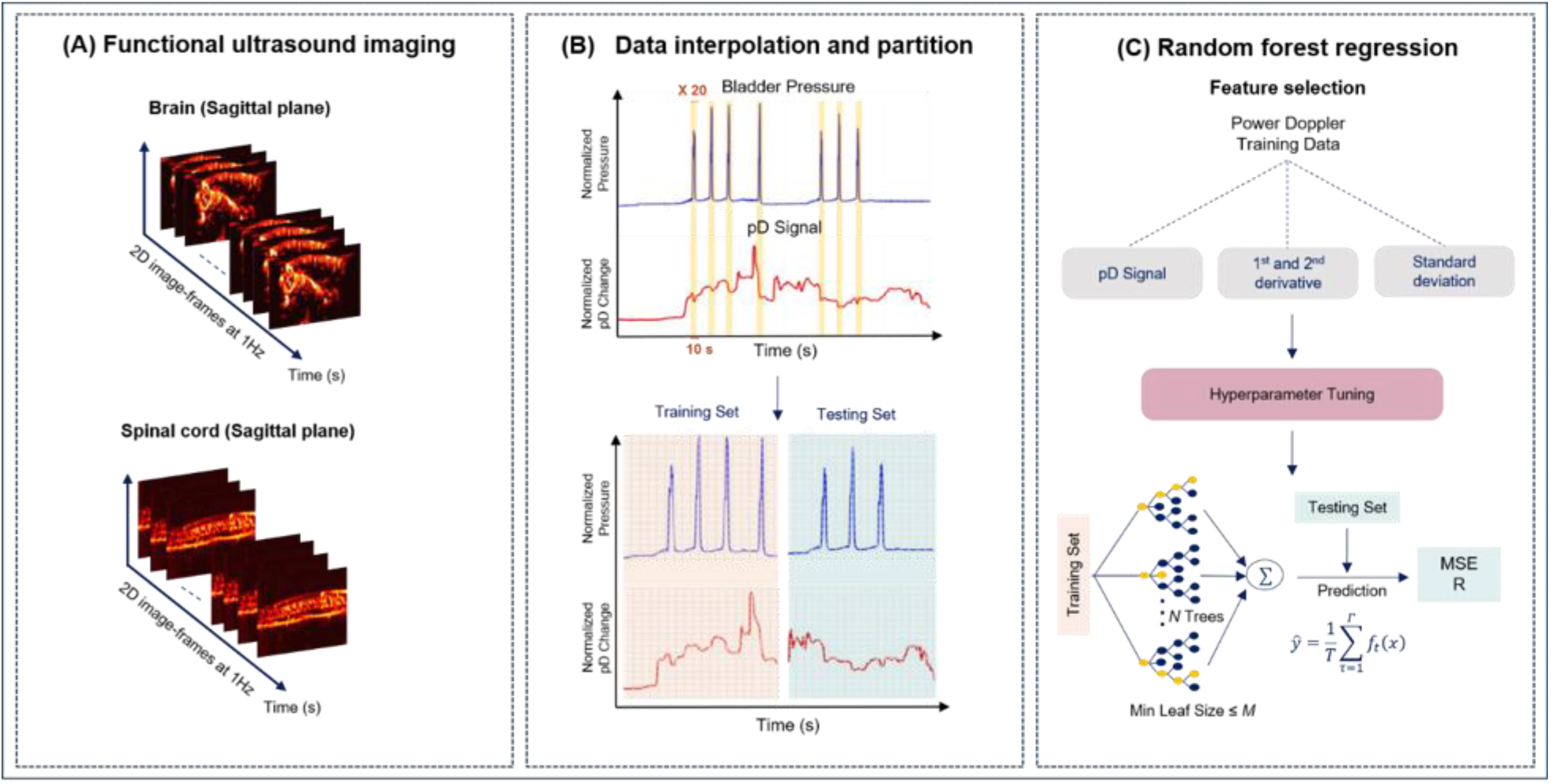
Random Forest Regression (RFR) framework for decoding bladder pressure from brain and spinal cord fUSI activity. **(A)** fUSI recordings acquired fromthe brain (n = 5 rats) or spinal cord (n = 7 rats) during simultaneous bladder filling and voiding experiments. **(B)** Preprocessing of normalized bladder pressure signals using piecewise cubic Hermite interpolation (PCHIP) with a factor of 20 within a ±5 s window around each voiding peak. The corresponding segments of the pD signal were extracted and interpolated as well. Data were then partitioned into a training set comprising the first half of the session (1–25 min; baseline, the first filling cycle, and the first half of the initial hold), whereas the second half (25–50 min; the remaining hold, the second filling cycle, and washout) was used to evaluate model performance. **(C)** Feature extraction from the training set, including pD signal, first and second derivatives of pD signal, and rolling standard deviation (using a backward-looking trailing window of 20 samples, to avoid future information leakage). Hyperparameters of the RFR model were optimized independently for brain and spinal cord datasets using grid search across ensemble size and minimum leaf size. Model performance was evaluated using mean squared error (MSE) to quantify prediction accuracy and Pearson correlation coefficient to assess correspondence between predicted and measured bladder pressure signals.

**Figures 6A** and **6B** illustrate the performance of the RFR model in predicting bladder pressure from brain and spinal cord pD signals, respectively, in two representative animals. In both cases, the predicted bladder pressure (orange traces) closely matched the actual bladder pressure dynamics (blue traces), capturing the increase during bladder filling and the rapid decrease associated with voiding. The model also exhibited high temporal precision in detecting voiding events, as reflected in the accurate timing of pressure transitions. Together, these findings indicate that the model effectively captured key features of the bladder cycle from brain and spinal cord recordings, reliably tracking pressure dynamics across filling and voiding phases. Model performance was quantified using Pearson correlation coefficient and mean squared error (MSE). For the brain group (n = 5), the mean Pearson correlation between actual and predicted bladder pressure was 0.5676 ± 0.049 (MEAN ± SE), and the corresponding MSE was 0.0273 ± 0.0045 (**Figure 6C**). For the spinal cord group (n = 7), the mean Pearson correlation was 0.7576 ± 0.051 (MEAN ± SE), and the corresponding MSE was 0.0138 ± 0.0017 (**Figure 6E**). Together, these results indicate that pD signals from both the brain and spinal cord encode physiologically relevant bladder pressure dynamics, enabling accurate and robust prediction of bladder pressure.

**Figure 6.**
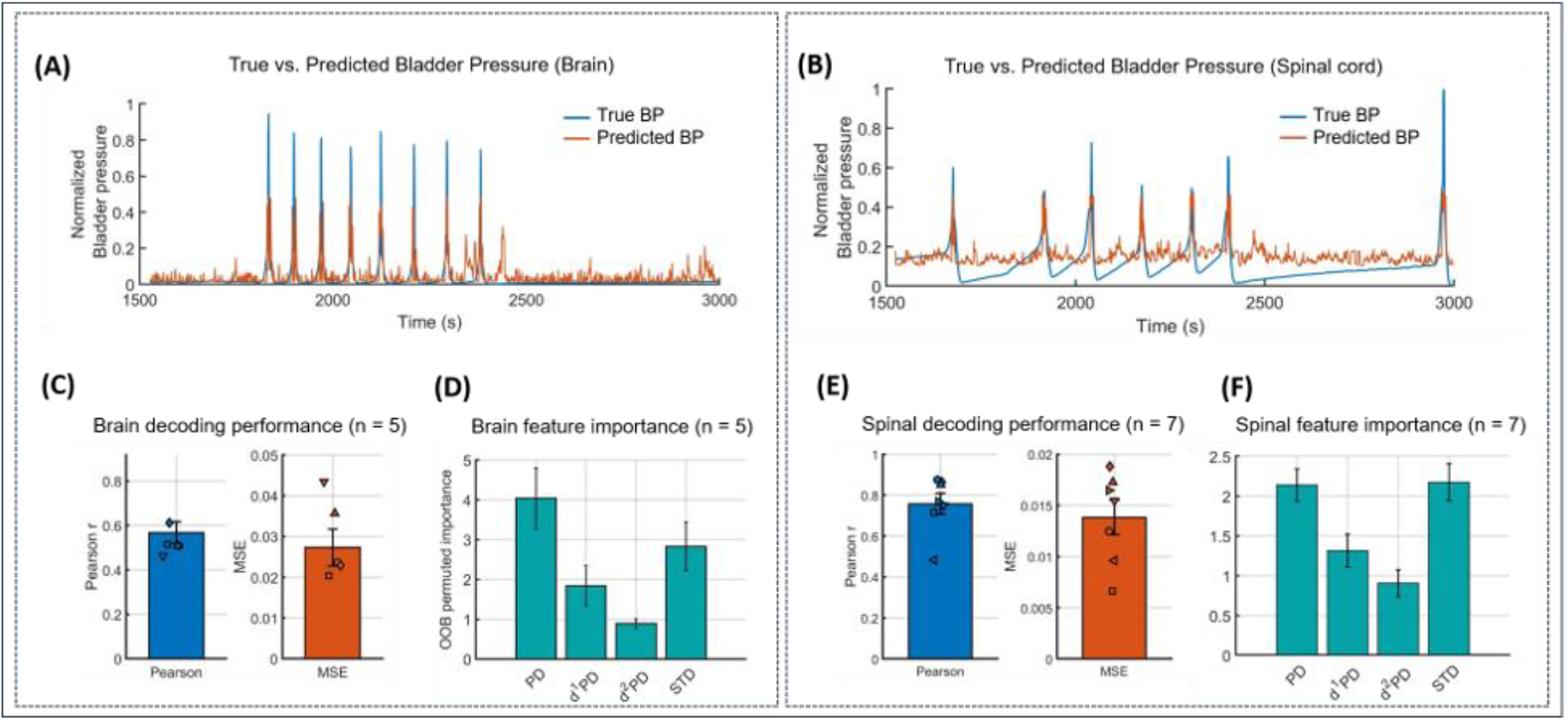
Random Forest Regression (RFR) decoding performance and feature contribution analysis for brain and spinal cord fUSI datasets. (A) Representative decoding performance for a rat in the brain group, showing predicted (orange) and true (blue) normalized bladder pressure traces. (B) Same as in panel A, but for a representative rat in the spinal cord group. (C) Decoding performance across animals in the brain group (n = 5), quantified using Pearson correlation coefficient and mean squared error (MSE) between predicted and measured bladder pressure signals. (D) Mean feature importance values across brain-group animals for each input feature, including the pD signal, first- and second-order temporal derivatives, and rolling standard deviation. Feature importance was estimated using permutation-based increases in out-of-bag prediction error. Values represent the mean increase in MSE following random permutation of each feature, normalized by the standard deviation of the error increase. Larger values indicate greater contribution to model performance. Individual markers correspond to values from single animals (E) Same as in panel C, but for the spinal cord group (n = 7). (F) Same as in panel D, but for spinal-cord group. Error bars in all panels (C-F) represent the standard error of the mean across animals.

To determine which pD-derived features contributed most strongly to model predictions, we computed average feature importance scores across animals. In the RFR model, feature importance reflects the relative contribution of each input feature to reducing prediction error across the ensemble of decision trees, with higher values indicating greater influence on model output. In the brain model (**Figure 6D**), the pD signal emerged as the most important feature (mean importance = 4.0406 ± 0.7617). In the spinal cord model, the rolling standard deviation showed the highest importance (2.1738 ± 0.2307), followed closely by the pD signal (2.1391 ± 0.2004), suggesting that both short-term variability in the pD signal and amplitude contributed strongly to prediction performance (**Figure 6F**). Prediction results for individual animals are provided in the Supplementary materials (**Figures S3 and S4**). Together, these findings demonstrate that pD signals from the brain and spinal cord reliably encode bladder dynamics, enabling robust decoding of pressure changes and accurate detection of spontaneous voiding events. These results establish hemodynamic signals as reliable indicators of bladder state across the micturition cycle.

## Discussion

### Overall

The current study reveals distributed neural dynamics across the brain and spinal cord associated with bladder filling and voiding. We identify robust and spatially distinct patterns of activation and deactivation across central nervous system structures. Furthermore, hemodynamic signals from the brain and spinal cord encode bladder pressure dynamics with sufficient fidelity to enable accurate prediction, supporting the development of brain-spinal cord machine interface technologies for patients with neurogenic bladder dysfunction.

### Neural substrates of bladder control in the brain

Our findings reveal a distributed brain network dynamically engaged during micturition, captured at high temporal resolution. By continuously imaging brain activity during bladder infusion that mimics physiological urine accumulation, we resolved how this distributed network engages and disengages around naturally elicited voiding reflexes. Activation spanned the sensorimotor and retrosplenial cortices, hippocampus, diencephalon, brainstem, and cerebellum, broadly consistent with current models of central bladder control^1,5,23^. In particular, the observed midbrain activity likely reflects engagement of regions encompassing the periaqueductal gray (PAG), a key integrative hub for bladder afferent input and voiding initiation^23–25^ – although the spatial resolution of our measurements does not permit precise delineation of specific nuclei. Importantly, our results extend these models by providing a spatiotemporally dynamic view of network activity across the peri-void period.

Several of the identified regions have previously been implicated in sensory, motor, cognitive, and autonomic aspects of bladder control^7,26–28^. Cortical regions are thought to contribute to voluntary control of voiding^9,29,30^, including pelvic floor contraction^10,11^, as well as processing bladder fullness and urinary urge^26^. Micturition is also strongly influenced by cognitive and affective processes, including attention, decision-making, emotional state and social context^1,31^, and dysfunction within these networks may contribute to neurogenic bladder disorders^32^. The retrosplenial cortex is considered analogous to the posterior cingulate cortex in humans and is involved in emotional processing and memory and maintains strong projections to the hippocampus^33,34^. Within this circuitry, the thalamus likely serves as a relay conveying signals from the PAG to the forebrain, including the cingulate cortex and insula^23,35,36^. Consistent with this integrative role, coherence analysis revealed strong coupling between bladder pressure and hemodynamic activity in the diencephalon, indicating continuous engagement of this region throughout the voiding cycle rather than responses restricted to discrete voiding events.

Consistent with prior studies, additional subcortical and cerebellar regions were also engaged during voiding^5,10,23^. Following urination onset, transient decreases in CBV emerged within subsets of brainstem and cerebellar regions, while most regions gradually returned toward pre-void levels by the end of voiding. In contrast, cerebellar activity remained elevated during the post-void period, suggesting sustained involvement beyond immediate voiding. This observation is consistent with the broader role of the cerebellum in autonomic regulation and sensorimotor coordination^37,38^.

### Neural substrates of bladder control in the spinal cord

The present results provide direct evidence of structured, segment-specific hemodynamic dynamics across the lumbosacral spinal cord during micturition. We observed a clear rostro-caudal progression of activity, with early modulation emerging in rostral segments during the pre-void phase, followed by a caudal shift and peak activation during and after voiding. This spatiotemporal organization aligns with the distributed nature of spinal circuits involved in bladder control, which span multiple segments and integrate both autonomic and somatic components^39,40^. Notably, these findings provide the first spatiotemporally resolved characterization of spinal cord hemodynamic activity during micturition, enabling direct visualization of segment-level dynamics that have not been accessible with prior neuroimaging approaches.

The spatial distribution of responses across the L2-S1 field of view is broadly consistent with known spinal cord organization. The parasympathetic preganglionic neurons extend into the lateral dorsal horn and dorsal commissure, regions that contain interneurons involved in lower urinary tract functions^1,41,42^. Bladder and urethral afferents terminate in overlapping regions of the lateral dorsal horn and the dorsal commissure^36,43^, supporting their role in viscerosomatic integration and coordination of micturition-related activity.

Our findings provide spatiotemporal functional evidence consistent with this organization. The positive hemodynamic responses observed in caudal-ventral regions during voiding are spatially consistent with the location of the dorsolateral motor neuron pool of the ventral horn – the rodent homolog of Onuf’s nucleus^44,45^ – and are consistent with the rhythmic bursting activity of the external urethral sphincter (EUS) motoneurons that drives urine expulsion in rats^46,47^. This is followed by a strong increase in SCBV in caudal-dorsal regions after voiding, overlapping with the L6-S1 termination zone of bladder and urethral afferents in the lateral dorsal horn and dorsal commissure. This signal may reflect afferent input from the bladder and urethra and downstream integration by dorsal horn interneurons^41,42,48^. In contrast, the early negative hemodynamic responses in rostro-ventral regions preceding voiding may indicate modulation of propriospinal coordinating circuits engaged in the bladder-EUS coordination required for void initiation^49^.

The observed spatiotemporal patterns further suggest coordinated interactions across spinal segments. An early decrease in hemodynamic signal in rostral regions was followed by increases in caudal segments, consistent with a transition from pre-void modulation to the execution of bladder emptying. This interpretation is supported by the ERA analysis, which revealed region-specific differences in both the magnitude and temporal evolution of responses, indicating functional heterogeneity along both the rostro-caudal and dorso-ventral axes.

Importantly, the multi-segmental and propagating nature of the activation patterns, spanning multiple contiguous lumbosacral levels, suggests that bladder control is not confined to a single spinal segment but involves coordinated recruitment across distributed spinal networks. Such coordinated activity is consistent with known descending and spinobulbospinal pathways involved in bladder control, including lateral pontine regions associated with the pontine storage center^1,50,51^. During voiding, the spinobulbospinal reflex pathway is conveyed through the pontine micturition center, which stimulates parasympathetic outflow to the bladder and urethral smooth muscle while inhibiting the sympathetic and pudendal outflow to the urethral outlet. Although brain and spinal cord recordings were acquired independently in the present study, our findings support the involvement of both supraspinal and spinal circuits in micturition.

Together, these findings extend existing anatomical and electrophysiological knowledge by providing spatiotemporally resolved functional evidence of how lumbosacral spinal circuits are dynamically engaged during micturition. Rather than reflecting static regional roles, these findings suggest a dynamic reorganization of spinal activity, characterized by shifting patterns of activation and deactivation across the micturition cycle.

### Decoding bladder pressure dynamics from the pD signal

Restoring bladder control is a top priority for paralyzed patients and is often ranked above restoration of motor and sensory functions. Similar to brain-machine interfaces for motor control, developing neural interfaces for bladder dysfunction requires first demonstrating reliable decoding of bladder pressure from neural signals. Here, we provide direct evidence that the pD signal recorded from the brain and spinal cord can be used to accurately decode the bladder pressure dynamics during the micturition cycle. To our knowledge, this is the first study to demonstrate that distributed brain and spinal cord activity can accurately predict bladder pressure changes.

Building on our previous work, which showed that spinal cord activity from a single spinal level could reconstruct bladder pressure during one urodynamically controlled filling and emptying cycle^21^, the present study extends these findings in several important ways. First, we demonstrate prediction rather than reconstruction by training and testing models across multiple micturition cycles, reducing the likelihood that decoding performance reflects overfitting to a single voiding event. Second, we show that bladder pressure can be predicted from distributed brain activity and multi-segmental spinal cord dynamics, providing a more comprehensive representation of the neural circuits underlying bladder control.

Unlike electrophysiological approaches, which often require the implantation of penetrating electrodes into neural tissue, our method offers a minimally invasive alternative with strong translational potential. While electrophysiology provides superior temporal resolution, chronic electrode implantation poses significant clinical challenges, including tissue damage, inflammation, and signal degradation over time. These risks are particularly pronounced in the spinal cord, where electrode placement carries a substantial danger of further neurological injury or paralysis, limiting feasibility for long-term or widespread use. Overall, these findings provide a first proof-of-concept that fUSI can be used for decoding internal organ states and advancing brain and spinal cord machine interface technologies for restoring bladder control.

### Limitations

Despite its high spatiotemporal resolution and sensitivity to hemodynamic changes, current 2D fUSI technology is limited to a single imaging plane. While this limits spatial coverage, emerging 3D fUSI approaches, enabled by next-generation ultrasound probes, are expected to substantially expand volumetric imaging of the brain and spinal cord^52,53^. In the present study, the imaging plane was positioned approximately along the midline to target medial midbrain structures. Consequently, more lateral regions, such as the insula and pallidum, which are implicated in interoception and the modulation of micturition^28,30,54^, were not captured within the field of view. This limitation is balanced by key advantages of the current approach, including system compactness and the feasibility of real-time decoding, which are critical for developing closed-loop brain and spinal cord machine interfaces^55^.

An additional consideration arises when imaging the spinal cord. Unlike the head, which can be rigidly secured in a stereotaxic frame, the spinal cord remains relatively mobile, introducing greater variability in imaging plane alignment across animals. Although the probe was consistently positioned in the midsagittal plane spanning the laminectomy window in all animals, and pD image registration was applied to align datasets to a common vascular reference across subjects, residual differences in positioning may have contributed to the increased variability observed in spinal cord activation patterns during micturition compared to the brain.

While urethane has been reported to attenuate EUS EMG activity^56^, we observed spontaneous voiding cycles throughout our recordings, alongside robust bladder-pressure-correlated neural activity in both the brain and spinal cord. Together, these findings indicate that the central micturition circuitry was actively engaged. The brain activity reported here therefore likely reflects reflex-related processing of bladder afferent input and central modulation of micturition, rather than voluntary or decision-related control, and represent a foundational layer of central bladder control that is preserved under anesthesia. Furthermore, we identified robust neural activity in both the brain and spinal cord that was strongly correlated with bladder pressure. Importantly, these signals enabled accurate prediction of bladder pressure dynamics, demonstrating that the underlying neural representations of bladder function are preserved and can be reliably decoded even under anesthetized conditions. On the other hand, although fUSI has been successfully applied in awake, behaving animals for brain imaging^57,58^, extending this approach to the spinal cord remains technically challenging and has not yet been demonstrated. Our findings therefore establish a critical foundation for future studies aimed at translating this approach to awake animals, where simultaneous brain and spinal cord imaging could further advance the development of brain-spinal cord machine interfaces for bladder control.

Finally, achieving accurate decoding performance with the RFR machine learning model requires high temporal resolution recordings of both pD signal and bladder pressure. Here, the temporal resolution of 1 s was increased by interpolating the signals by a factor of 20 around the voiding peak, a step necessary to augment the number of training samples during the relative brief phase of rapid bladder pressure changes associated with voiding. Future studies will aim to directly acquire higher-frame-rate pD signal and bladder pressure data. Notably, fUSI systems are capable of temporal sampling on the order of tens of milliseconds^59^. Although higher frame rates could improve decoding accuracy by capturing faster signal dynamics, they come at the cost of reduced pD sensitivity. This trade-off would have limited our ability to detect subtle, spatially distributed hemodynamic changes, which are critical for identifying brain and spinal cord regions involved in micturition. Power Doppler sensitivity depends on integrating signals across an ensemble of ultrafast frames, increasing the temporal sampling rate shortens this ensemble, weakening clutter rejection and reducing signal-to-noise – an effect that disproportionately compromises the detection of small responsive regions^60^. In future work, when decoding rather than spatial mapping is the primary objective, fUSI acquisition parameters can be optimized for higher temporal resolution to better support real-time decoding applications.

## Conclusions

In summary, we identify distributed neural substrates across the brain and spinal cord associated with micturition and show that this activity encodes bladder pressure dynamics that can be accurately decoded using fUSI. By linking spatially resolved neural responses to bladder function, this work establishes a foundation for brain and spinal cord machine interface approaches to restore bladder control in patients with neurogenic bladder dysfunction - a technology that currently does not exist.

## Methods

### Animals

Five (5) and seven (7) female Sprague-Dawley rats (250 - 350 g, Charles River Laboratories; Hollister, CA) were used for brain and spinal cord imaging, respectively. Rats were acclimatized for at least 7-days prior to surgery. All procedures were approved by the University of Southern California, Institutional Animal Care and Use Committee (IACUC #21319).

#### Anesthesia

Anesthesia was induced with 5% isoflurane, followed by subcutaneous administration of urethane at a dose of 1.4 g/kg. Body temperature of the animals was maintained using an underbody warming pad. Adequate depth of anesthesia was monitored by the absence of reflex responses (e.g., toe-pinch reflex) and by monitoring respiratory rate.

### Bladder catheterization

An anterior midline suprapubic incision was performed, and the abdominal wall was dissected in layers to access the peritoneal cavity and to expose the urinary bladder. The bladder dome was punctured using an 18-gauge needle, through which a pressure transducer and urinary catheter (Millar Instruments, Texas, US) were inserted into the bladder^61^ and secured in a watertight fashion using an 8-0 braided purse-string suture. The abdominal wall was subsequently closed using nylon suture, and animals were turned into prone position for spinal or cranial surgery (see following sections). Throughout experiments, the bladder pressure was continuously measured using a pressure transducer (Millar Instruments, Texas, US) and recorded with PowerLab 4/35 data acquisition system (ADInstruments, Colorado, US) at a sampling rate of 1 kS/s. The experimental set-up is illustrated in **Figure 1A**.

### Craniectomy

Five (5) rats were included in the intracranial recording portion of this study. The head was fixed in a stereotactic frame using ear-bars (Kopf Instruments, California, US) to minimize motion artifact. The scalp hair was shaved and further chemically removed using a depilatory cream (Nair; Church & Dwight Co., Inc, New Jersey, US). A midline scalp incision was then made, extending from the level of the orbit to the upper cervical spine. The periosteum was scraped off, and a craniectomy was performed using a handheld electric drill. The craniectomy was performed with the anterior border at bregma, posterior border just rostral to the foramen magnum (total anteroposterior extent spanning approximately 10 mm), and lateral borders at the superior temporal line (width spanning approximately 8 mm). The craniectomy was sized to accommodate the ultrasound probe and to provide coverage of the frontal cortex, diencephalon, midbrain, and rostral cerebellum (**Figure 1B**). Hemostasis was achieved by gentle tamponade of bridging veins draining into venous sinuses. Intraoperative blood losses were compensated with 1-2 ml intramuscular injections of 0.9% saline.

### Laminectomy

Seven (7) rats were included in the spinal cord recording portion of the study. Heads of animals were fixed in the stereotaxic frame, and a laminectomy spanning the T12-L1 vertebrae was performed to expose spinal cord levels of L2-S1 (**Figure 1C**). The ultrasound probe was positioned in a mid-sagittal plane to enable simultaneous imaging of multiple spinal cord levels. Note that axial fUSI acquisition of the spinal cord was not performed, as prior work^62^ in rats indicates that it is suboptimal due to noise from surrounding bones and muscles, particularly given the small cross-sectional area of the rat spinal cord.

### Urodynamic measurements

A simultaneous urodynamics and imaging protocol consisting of 10-min baseline period, followed by two cycles of continuous bladder filling (10 min) interspersed by a hold period (10 min), and completed with additional washout (10 min) period acquisitions was used for the study (**Figure 1D**). Bladder filling was performed using a micro-injector pump (Chemyx, Stafford, Texas) to infuse 0.9% saline at a constant rate of 0.1 ml/min for 10 min. Throughout the experiment, including both filling, hold and washout periods, animals were allowed to void spontaneously. A representative bladder pressure trace in **Figure 1E** shows void peaks (red dots) during filling periods.

### Functional ultrasound imaging

We used an Iconeus One scanner (Iconeus, Paris, France) to acquire pD images from the brain and spinal cord of the animals. The scanner was equipped with a miniaturized 128 elements linear array probe, with 15 MHz center frequency, 0.1 mm pitch that generates pD images with a spatial resolution of 100 μm × 100 μm in-plane, slice thickness of 400 μm and field of view (FOV) 9.1 mm × 12.8 mm (depth and width). Each pD image was obtained from 200 compounded frames acquired at 500 Hz frame rate, using 11 tilted plane waves offset by 2° (i.e., from -10° to +10° increment by 2°) acquired at a 5500 Hz pulse repetition frequency. Successive blocks of 400 ms compounded plane wave imaging with 600 ms interval between blocks were used, resulting in a final frame rate of 1 image per second. The pD images were preprocessed within the Iconeus system using phase-correlation based sub-pixel motion registration^63^ and singular value decomposition (SVD)-based clutter filtering algorithms^64^ to separate tissue motion from hemodynamic signals^60^. The resulting pD images were then extracted for subsequent signal processing. **Figure 2A** and **2B** present pD vascular maps of the brain and spinal cord, respectively, from representative animals with vascular and anatomical annotation overlays. In both experiments, the ultrasound probe was mounted on and positioned using a three-axis motorized system. For brain imaging, a parasagittal plane located 1 mm to the right of the midline was acquired within the craniectomy window as described above. For spinal cord imaging, the probe was positioned in the midsagittal plane, spanning the laminectomy window, as described above.

### Data Preprocessing

To reduce physiological and motion-related artifacts in the pD images, we implemented piecewise rigid motion correction^65^, previously used and validated in fUSI^21,55,66,67^ and other neuroimaging studies^68–70^. To enhance signal stability, the motion correction algorithm was integrated with in-house developed respiratory filtering, high-frequency smoothing, and image stabilization. We applied a low-pass filter with a 0.04 Hz normalized passband and 60 dB stopband attenuation, which compensated for the phase delay introduced by the filter, to remove high-frequency fluctuations in the pD signals.

### pD Image Registration

An important next step is the accurate registration of pD vascular maps across animals to a common anatomical reference. Robust inter-animal registration is essential for pixel-based analyses and for extending fUSI toward more quantitative, group-level investigations. Such alignment allows the application of advanced computational techniques, including machine learning and statistical parametric mapping (SPM), that have been widely adopted in other neuroimaging modalities^71–73^. To establish a reliable reference pD image, a representative animal was selected from each group (brain and spinal cord) based on the clarity and anatomical fidelity of its pD vascular (**Supplementary Figures S1 and S2**). Distinct registration strategies were then applied: an affine transformation (scaling, rotation and shearing), was used for brain pD images to correct global geometric differences^74,75^, whereas a locally weighted mean (LWM) transformation (a polynomial warping based on selected neighboring control points), was applied to spinal cord pD images to accommodate regional shape variability^76^.

### Brain pD Image Registration

We registered the brain pD images (rats 1-5) to the reference animal (rat 1) using an affine transformation in MATLAB R2024b (MathWorks, Natick, MA). The affine transformation provides a linear mapping that corrects global geometric differences arising from minor variations in imaging orientation or field of view across animals, while preserving relative spatial relationships between anatomical structures^74,77^.

Transformation parameters were estimated using manually selected anatomical control points distributed across cortical and subcortical regions (*fitgeotrans*). The control points correspond to identifiable anatomical landmarks present in both the moving and reference images and served as anchors for estimating the spatial transformation needed to align the two images. The affine model used these landmark correspondences to compute translation, rotation, scaling, and shear parameters. The calculated transformation ensured spatial alignment between each animal’s brain pD image and the reference brain image. This resulting transformation was applied to the entire pD time series of each animal, while preserving pixel intensities. The resulting global, rigid transformation preserved the overall anatomical structure while compensating for minor positional and angular variations between datasets, thereby enabling cross-subject averaging without geometric distortion. **Supplementary Figure S1** visualizes the registration results for the brain pD images, showing individual pD images before and after registration overlayed onto the reference brain to assess alignment quality.

### Spinal Cord Registration

For spinal cord imaging (rats 6-12), we performed non-rigid nonlinear registration to align all spinal cord pD images to a reference animal (rat 8) using locally weighted mean (LWM) transformation^76^. The elongated geometry of the spinal cord and its inherent morphological variability, including curvature and minor torsion across animals, render global transformations insufficient for achieving accurate anatomical correspondence. The LWM approach employs local polynomial interpolation^76,78^ around manually selected control points, ensuring accurate alignment despite inherent curvature and shape variability among spinal cords, and thereby ensures reliable inter-subject registration. As in the brain registration procedure, the resulting transformation was applied to the full pD time series of each animal while preserving pixel intensity values. This nonlinear registration effectively corrected inter-animal shape differences while maintaining anatomical integrity. **Supplementary Figure S2** illustrates representative pD images before and after registration overlaid on the reference spinal cord to assess alignment accuracy.

### Quantify registration performance

To quantify the registration performance, we employed normalized cross-correlation (NCC), a widely used similarity metric in image processing. NCC measures the similarity between two images by computing the correlation coefficient between their corresponding intensity values, while it remains robust to linear changes in brightness and contrast^79^. The registration results demonstrated a marked improvement in spatial correspondence of pD images across animals. For the brain, the mean NCC increased from 0.4287 ± 0.030 (MEAN ± SE) before registration to 0.6196 ± 0.018 after registration. Similarly, for the spinal cord, the mean NCC improved from 0.4079 ± 0.059 (MEAN ± SE) to 0.6235 ± 0.020 following registration, indicating enhanced spatial alignment while preserving the geometric integrity of the original vascular structures.

### Coherence analysis

We performed coherence analysis to identify brain and spinal cord regions with pD signals exhibiting similar fluctuation patterns to bladder pressure. This analysis considers the entire recording duration and quantifies, for each pixel, the degree of synchrony between pD and bladder pressure signals across frequencies, independent of phase offsets. Coherence is analogous to Pearson’s correlation in the frequency domain and quantifies the degree of linear coupling between two signals at each frequency on a scale from 0 to 1, where 0 indicates no linear relationship and 1 indicates perfect linear dependence (i.e., one signal can be linearly predicted from the other at that frequency). As a useful statistic tool, coherence has been previously applied successfully to other functional imaging data^80,81^.

We computed magnitude-squared coherence (MSC) between each pD image-pixel signal and the bladder pressure trace (simultaneously recorded) to quantify frequency-specific coupling between hemodynamic activity and bladder pressure. All analyses were implemented in *MATLAB* using the built-in *mscohere* function. Coherence estimates were obtained using Welch’s averaged periodogram method ^82^, in which Fast Fourier Transforms (FFTs) were computed on overlapping, windowed segments of each time series to estimate power and cross-spectral densities (Hamming windows of 640 samples with 320-sample overlaps).

The coherence values were averaged within a frequency range (0.0033-0.033 Hz) corresponding to periods of 30 s to 5 min. This range encompasses the inter-void intervals while filtering out high-frequency components and physiological noise (e.g., cardiac and respiratory fluctuations). To visualize coherence patterns across animals, we computed group-averaged coherence maps separately for the brain and spinal cord. This produced a single mean coherence map per group that represents the average linear coupling between bladder pressure and pD signals within this frequency range.

### Dynamic statistical parametric mapping (dSPM)

While coherence analysis identifies regions whose hemodynamic activity is temporally coupled to bladder pressure dynamics, dynamic statistical parametric mapping (dSPM) characterizes the spatiotemporal evolution of brain and spinal cord activity before, during, and after voiding events. Specifically, dSPM was used to visualize transient changes in cerebral (ΔCBV) and spinal cord blood volume (ΔSCBV) associated with different phases of the voiding cycle.

Bladder pressure peaks corresponding to voiding events were identified in each animal based on the abrupt pressure drop following each peak (**Figure 1E).** For each event, pD signals were extracted within a ±16 s window centered on the peak, ensuring temporal isolation of individual voiding events. Peri-void bladder pressure and pD signals were averaged across events within each animal and subsequently across animals within each group (brain and spinal cord), yielding group-level time-resolved pD matrices (space × time). The resulting pD signal matrices were used for the dSPM analysis to examine the spatiotemporal evolution of voiding-related hemodynamic activity.

To generate the dSPM maps, percentage changes in the pD signal were computed within non-overlapping 3-s windows spanning −16 to +16 s relative to the voiding peak. A fixed pre-void baseline period was used as reference, defined as the mean pD signal during the first 3-s window (i.e., −16 to −14 s) of the peri-void interval. For each pixel and time window, z-scores were computed from the percentage signal changes relative to this reference activity. Statistical significance of voiding-related changes was assessed using a two-sided Student’s *t*-test comparing each time window to the reference period. To account for multiple pixel-wise comparisons, false discovery rate (FDR) correction was applied using the Benjamini–Hochberg procedure with a significance threshold of p < 0.01^83,84^. This approach enables identification of temporally specific and spatially distributed patterns of activation and deactivation during transition from pre-to post-voiding periods.

Activation maps were generated to identify regions exhibiting positive and negative responses to bladder pressure changes. For visualization of dSPM maps, all significantly modulated pixels were retained, whereas low magnitude z-scores were rendered transparent to reduce visual crowding, thereby highlighting the spatial distribution of stronger positive and negative hemodynamic responses. A Gaussian smoothing kernel (σ = 1) was applied for display.

Finally, to quantify the time-varying spatial evolution of voiding-related hemodynamic changes, we defined a pixel-count threshold based on the top 15% of significantly modulated pixels within the window surrounding void peak (−1 to +1), and applied this threshold to brain and spinal cord dSPMs across successive 3-s time windows to track the spatial extent of hemodynamic activity over time. This threshold corresponded to z-score cutoffs of (z < −1.35 or z > +1.582) for the brain and (z < −1.488 or z > +1.537) for the spinal cord. This analysis enabled tracking of temporal fluctuations in the spatial distribution of activation and deactivation among the most significantly modulated pixels.

To complement the dSPM analysis of spatiotemporal activation patterns, event-related average (ERA) analysis was used to elucidate the temporal profile of hemodynamic responses within brain and spinal cord ROIs, expressed as percentage changes in CBV (ΔCBV) and SCBV (ΔSCBV) relative to the 3-s reference period (−16 to −14 s). The dSPM frame centered on the voiding peak (−1 to +1 s) was used to quantify the percent pD signal changes within each ROI based on the significantly activated pixels, and the mean positive and negative response for each ROI was then plotted as ERAs. To compare peak responses across ROIs, we conducted pairwise comparisons of peak amplitudes using a nonparametric approach. Analyses were performed separately for positively and negatively activated pixels. For each ROI, peak values were extracted and compared across all ROI pairs using the two-sided Wilcoxon rank-sum tests, thereby avoiding assumptions of normality in the distribution of peak responses. Only ROI pairs in which both regions contained at least five significantly activated pixels were included to ensure sufficient sample size for reliable nonparametric comparison. Resulting p-values were corrected for multiple comparisons using the Benjamini – Hochberg false discovery rate (FDR) procedure (α < 0.05) ^83,84^.

### Decoding of bladder pressure changes from brain and spinal cord activity

We implemented a random forest regression model (RFR) using a bagged ensemble of regression trees to assess whether pD signals recorded from the brain and spinal cord can decode (i.e., predict) bladder pressure changes **(Figure 5)**. This nonparametric ensemble learning approach combines the outputs of multiple decision trees, each trained on a bootstrap-resampled subset of the training data. By averaging predictions across trees, bagging reduces model variance, limits overfitting, and improves generalization ^85,86^. We chose this approach because tree-based ensembles are ideally suited to handle multicollinearity, capture complex interactions without explicit specification, and maintain high accuracy with relatively small sample sizes^87,88^.

Decoding bladder pressure from the pD signal involves five parts: (a) interpolation of bladder pressure and pD signals around voiding events to enhance temporal resolution, (b) extraction of features from the pD signal, (c) development and training of the RFR model, (d) evaluation of model performance using statistical metrics, and (e) comparison and visualization of predicted versus observed bladder pressure traces. Interpolation was necessary because voiding is a rapid process characterized by abrupt changes in bladder pressure, and the fUSI data were acquired at a relative low temporal resolution (1 Hz). Although the fUSI scanners are capable of substantially higher temporal sampling (down to ∼10 ms), the acquisition sequence employed in this study prioritized spatial resolution and sensitivity over temporal resolution. To enhance temporal resolution around voiding events, we applied piecewise cubic Hermite interpolation (PCHIP), which preserves the shape and monotonicity of the original signal^89^. We selected a ±5 second window centered on each voiding peak and interpolated the data within this window by factor of 20, yielding a denser and more uniform temporal sampling around voiding events. Prior to interpolation, bladder pressure and pD signals were normalized to the range [0, 1] to ensure consistent scaling across datasets and to facilitate effective training and convergence of the machine learning model. We then smoothed the bladder pressure signals using a Gaussian filter to reduce high-frequency noise while preserving overall signal dynamics **(Figure 1E, blue curve)**. To ensure reliable performance evaluation and avoid overfitting, we employed a cross-validation strategy where the first half of the session (1-25 min; baseline, first filling cycle, and the first half of the initial hold) was used for model training, and the second half (25-50 min; remaining hold, second filling cycle, and washout) was used to test the model performance. We deliberately avoided random splitting of the dataset, as such a strategy could result in temporally adjacent data points being assigned to both training and test sets. Given the strong temporal autocorrelation in the pD signal, this could artificially inflate model performance by introducing information leakage. By partitioning the dataset along well-defined temporal boundaries, we ensured a more robust and realistic assessment of the model’s predictive capacity.

Feature extraction was performed on the training set by constructing a feature matrix from the preprocessed pD signal and its derived features: (1) the first derivative, capturing temporal trends and rates of change; (2) the second derivative, reflecting acceleration-like dynamics and inflection points; and (3) the rolling standard deviation, quantifying local signal variability over time and computed using a trailing window of 20 samples (i.e., at each time point, the standard deviation was calculated from the current and 19 preceding samples). This trailing-window approach ensured that only past information was used at each time point, thereby preventing information leakage from future samples. The rolling standard deviation enabled the model to capture local signal fluctuations that may reflect physiologically relevant transitions, such as voiding onset or changes in bladder pressure buildup. Together, the features provided a comprehensive representation of the temporal dynamics of the pD signal and improved the ability of the model to capture underlying bladder pressure changes.

To develop the RFR model, we performed a systematic grid search across all animals to optimize hyperparameters governing tree complexity and ensemble size. Specifically, we evaluated the influence of the minimum leaf size of individual regression trees and the number of learning cycles in the bagged ensemble on model performance. Model hyperparameters were tuned over minimum leaf size [10, 25, 50, 100, 250, 500, 750, 1000] samples and number of learning cycles [50, 100, 250, 500, 750, 1000, 1500, 2000]. For each hyperparameter combination, a bagged ensemble of regression trees was trained on the training segment and subsequently evaluated on a held-out test segment, with minimum leaf size restricted to values smaller than the corresponding training-set size. Performance was then quantified using Pearson correlation and mean squared error (MSE). To obtain unbiased group-level average, correlation coefficients were Fisher z-transformed prior to averaging across animals and subsequently back-transformed for reporting. In addition to the lowest mean test error, we also examined the variability of performance across animals and the train–test error gap to identify a model that generalized robustly. Model performance was evaluated as a function of the number of trees and minimum leaf size to identify stable, high-performing regions of hyperparameter space, where generalization performance was both high and stable across animals.

We implemented the RFR model using the MATLAB *fitrensemble* function, with bootstrap aggregation (bagging) of decision trees^85,90^. Using the hyperparameters identified through optimization, we trained separate models for the brain and spinal cord groups to predict bladder pressure, with the brain model using *100* trees and a minimum leaf size of *50*, and the spinal cord model using *500* trees and a minimum leaf size of *100*. These parameter choices were selected to mitigate overfitting while capturing non-linear relationships in the data. The final model prediction *f̂bag(x)* was computed by averaging the outputs of all B trees: Where *T_b_*_(*x*)_ denotes the output of the *b-th* tree and *B* s the total number of trees. Each tree is trained using a bootstrap sample of the training set, a process known as “bagging”, and considers only a random subset of predictors at each split ^91^. This makes RFR particularly suitable for modeling physiological signals with nonlinear and interactive effects.

To assess the contribution of individual predictors, we computed feature importance scores using the *out-of-bag (OOB) permutation method* implemented in MATLAB. Specifically, we used the *oobPermutedPredictorImportance* function, which quantifies the importance of each feature by measuring the increase in prediction error when the values of that feature are randomly permuted in the OOB samples. This method identifies pD-derived features that contribute most strongly to bladder pressure prediction while accounting for feature interactions and model complexity. The *Importance* score for each feature *j* was computed as:

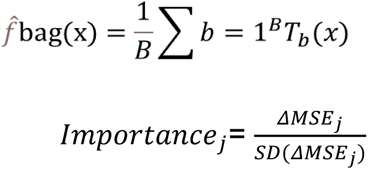

where *ΔMSE_j_* represents the change in OOB mean squared error after permuting feature *j*, and *SD*(*ΔMSE_j_*) is the standard deviation of that change across all trees. Higher importance scores indicate greater predictive contribution of the corresponding feature^85^. Such permutation-based importance metrics are widely used in RFR modeling due to their robustness and interpretability^90,92^.

### Software analysis

All data pre- and post-processing and statistical analysis were performed using Matlab Version (R2024b, R2025b).

## Data availability

The datasets generated and analyzed during the current study are available from the corresponding authors on reasonable requests. The datasets cannot be used for commercial purposes.

## Acknowledgments

This work was supported by the “Brain Research Foundation (BRF)”, “The USC Neurorestoration Center” at the University of Southern California and the “Alfred E. Mann Department of Biomedical Engineering” of the University of Southern California through start-up funding.

## Author Contributions

V.R.E, E.K, D.J.L, C.Y.L and V.C conceived the study. Y.T.L, K.W, H.Z, W.C and A.A performed the surgeries and acquired the data. H.F.Z, S.S, K.A.A and Y.T.L performed the data processing and analysis. H.F.Z, S.S and Y.T.L drafted the manuscript with substantial contributions from K.A.A, D.J.L, E.K, C.Y.L and V.C. All authors edited and approved the final version of the manuscript. C.L. and V.N.C. supervised the research.

## References

1. Fowler, C. J., Griffiths, D. & Groat, W. C. De. The neural control of micturition. Nat. Rev. Neurosci. 9, 453–466 (2008).

2. Kuhtz-Buschbeck, J. P. et al. Control of bladder sensations: An fMRI study of brain activity and effective connectivity. Neuroimage 47, 18–27 (2009).

3. Fukuyama, H. et al. Neural control of micturition in man examined with single photon emission computed tomography using 99mTc-HMPAO. Neuroreport 7, 3009–3012 (1996).

4. Blok, B. F. M., Sturms, L. M. & Holstege, G. Brain activation during micturition in women. Brain 121 (Pt 11), 2033–2042 (1998).

5. Blok, B. F. M., Willemsen, A. T. M. & Holstege, G. A PET study on brain control of micturition in humans. Brain 120 (Pt 1), 111–121 (1997).

6. Pang, D., Gao, Y., Liao, L. & Ying, X. Brain functional network alterations caused by a strong desire to void in healthy adults: a graph theory analysis study. Neurourol. Urodyn. 39, 1966–1976 (2020).

7. Griffiths, D., Derbyshire, S., Stenger, A. & Resnick, N. Brain control of normal and overactive bladder. J. Urol. 174, 1862–1867 (2005).

8. Matsumoto, S., Ishikawa, A., Matsumoto, S. & Homma, Y. Brain response provoked by different bladder volumes: a near infrared spectroscopy study. Neurourol. Urodyn. 30, 529–535 (2011).

9. Kuhtz-Buschbeck, J. P. et al. Cortical representation of the urge to void: a functional magnetic resonance imaging study. J. Urol. 174, 1477–1481 (2005).

10. Zhang, H. et al. An fMRI study of the role of suprapontine brain structures in the voluntary voiding control induced by pelvic floor contraction. Neuroimage 24, 174–180 (2005).

11. Seseke, S. et al. Voluntary pelvic floor muscle control-an fMRI study. Neuroimage 31, 1399–1407 (2006).

12. Pang, D., Gao, Y. & Liao, L. Functional brain imaging and central control of the bladder in health and disease. Front. Physiol. 13, (2022).

13. Giove, F. et al. Issues about the fMRI of the human spinal cord. Magn. Reson. Imaging 22, 1505–1516 (2004).

14. Powers, J. M., Ioachim, G. & Stroman, P. W. Ten Key Insights into the Use of Spinal Cord fMRI. Brain Sci. 8, (2018).

15. Stroman, P. W. et al. The current state-of-the-art of spinal cord imaging: Methods. Neuroimage 84, 1070–1081 (2014).

16. Eippert, F., Kong, Y., Jenkinson, M., Tracey, I. & Brooks, J. C. W. Denoising spinal cord fMRI data: Approaches to acquisition and analysis. Neuroimage 154, 255–266 (2017).

17. MacÉ, E. et al. Functional ultrasound imaging of the brain. Nat. Methods 8, 662– 664 (2011).

18. Tanter, M. & Fink, M. Ultrafast imaging in biomedical ultrasound. IEEE Trans. Ultrason. Ferroelectr. Freq. Control 61, 102–119 (2014).

19. Bercoff, J. et al. Ultrafast compound Doppler imaging: providing full blood flow characterization. IEEE Trans. Ultrason. Ferroelectr. Freq. Control 58, 134–147 (2011).

20. Mace, E. et al. Functional ultrasound imaging of the brain: theory and basic principles. IEEE Trans. Ultrason. Ferroelectr. Freq. Control 60, 492–506 (2013).

21. Agyeman, K. A. et al. Human spinal cord activation during filling and emptying of the bladder. Nature Communications 2025 16:1 16, 6506 (2025).

22. Paxinos, George. & Watson, Charles. The rat brain in stereotaxic coordinates. (2005).

23. Tai, C. et al. Brain switch for reflex micturition control detected by FMRI in rats. J. Neurophysiol. 102, 2719–2730 (2009).

24. Griffiths, D. Neural control of micturition in humans: a working model. Nature Reviews Urology 2015 12:12 12, 695–705 (2015).

25. Blok, B. F. M. & Holstege, G. Direct projections from the periaqueductal gray to the pontine micturition center (M-region). An anterograde and retrograde tracing study in the cat. Neurosci. Lett. 166, 93–96 (1994).

26. Athwal, B. S. et al. Brain responses to changes in bladder volume and urge to void in healthy men. Brain 124, 369–377 (2001).

27. Griffiths, D., Tadic, S. D., Schaefer, W. & Resnick, N. M. Cerebral control of the bladder in normal and urge-incontinent women. Neuroimage 37, 1–7 (2007).

28. Nour, S., Svarer, C., Kristensen, J. K. I., Paulson, O. B. & Law, I. Cerebral activation during micturition in normal men. Brain 123 (Pt 4), 781–789 (2000).

29. Fowler, C. J. & Griffiths, D. J. A decade of functional brain imaging applied to bladder control. Neurourol. Urodyn. 29, 49–55 (2010).

30. Griffiths, D. J. & Tadic, S. D. Bladder control, urgency, and urge incontinence: evidence from functional brain imaging. Neurourol. Urodyn. 27, 466–474 (2008).

31. Wood, S. K., Baez, M. A., Bhatnagar, S. & Valentino, R. J. Social stress-induced bladder dysfunction: potential role of corticotropin-releasing factor. Am. J. Physiol. Regul. Integr. Comp. Physiol. 296, R1671 (2009).

32. Fowler, C. J. Neurological disorders of micturition and their treatment. Brain 122 (Pt 7), 1213–1231 (1999).

33. Wyass, J. M. & Groen, T. Van. Connections between the retrosplenial cortex and the hippocampal formation in the rat: a review. Hippocampus 2, 1–11 (1992).

34. Maddock, R. J. The retrosplenial cortex and emotion: New insights from functional neuroimaging of the human brain. Trends Neurosci. 22, 310–316 (1999).

35. Tang, P. C. & Ruch, T. C. Localization of brain stem and diencephalic areas controlling the micturation reflex. J. Comp. Neurol. 106, 213–245 (1956).

36. de Groat, W. C., Griffiths, D. & Yoshimura, N. Neural control of the lower urinary tract. Compr. Physiol. 5, 327–396 (2015).

37. Paulin, M. G. The Role of the Cerebellum in Motor Control and Perception. Brain Behav. Evol. 41, 39–50 (1993).

38. Romano, V. et al. Functional Convergence of Autonomic and Sensorimotor Processing in the Lateral Cerebellum. Cell Rep. 32, 107867 (2020).

39. Yu, D. Direct projection from the lumbosacral spinal cord to spinally-projecting neurons in Barrington’s nucleus: an electron microscopic study in the rat. (2002).

40. Birder, L. A., Roppolo, J. R., Erickson, V. L. & De Groat, W. C. Increased c-fos expression in spinal lumbosacral projection neurons and preganglionic neurons after irritation of the lower urinary tract in the rat. Brain Res. 834, 55–65 (1999).

41. Groat, W. C. De, Vizzard, M. A., Araki, I. & Roppolo, J. Spinal interneurons and preganglionic neurons in sacral autonomic reflex pathways. Prog. Brain Res. 107, 97–111 (1996).

42. Nadelhaft, I. & Booth, A. M. The location and morphology of preganglionic neurons and the distribution of visceral afferents from the rat pelvic nerve: a horseradish peroxidase study. J. Comp. Neurol. 226, 238–245 (1984).

43. Thor, K. B., Morgan, C., Nadelhaft, I., Houston, M. & de Groat, W. C. Organization of afferent and efferent pathways in the pudendal nerve of the female cat. J. Comp. Neurol. 288, 263–279 (1989).

44. McKenna, K. E. & Nadelhaft, I. The organization of the pudendal nerve in the male and female rat. Journal of Comparative Neurology 248, 532–549 (1986).

45. Koliatsos, V. E., Price, D. L. & Clatterbuck, R. E. Motor neurons in Onuf’s nucleus and its rat homologues express the p75 nerve growth factor receptor: sexual dimorphism and regulation by axotomy. J. Comp. Neurol. 345, 510–527 (1994).

46. Kruse, M. N., Noto, H., Roppolo, J. R. & de Groat, W. C. Pontine control of the urinary bladder and external urethral sphincter in the rat. Brain Res. 532, 182– 190 (1990).

47. Peng, C. W., Chen, J. J. J., Cheng, C. L. & Grill, W. M. Role of pudendal afferents in voiding efficiency in the rat. Am. J. Physiol. Regul. Integr. Comp. Physiol. 294, (2008).

48. Fuller-Jackson, J. P., Osborne, P. B. & Keast, J. R. Regional Targeting of Bladder and Urethra Afferents in the Lumbosacral Spinal Cord of Male and Female Rats: A Multiscale Analysis. eNeuro 8, (2021).

49. Karnup, S. V. & de Groat, W. C. Propriospinal Neurons of L3-L4 Segments Involved in Control of the Rat External Urethral Sphincter. Neuroscience 425, 12–28 (2020).

50. Griffiths, D. J. The pontine micturition centres. Scand. J. Urol. Nephrol. Suppl. 36, 21–26 (2002).

51. Holstege, G., Griffiths, D., Wall, H. De & Dalm, E. Anatomical and physiological observations on supraspinal control of bladder and urethral sphincter muscles in the cat. J. Comp. Neurol. 250, 449–461 (1986).

52. Bertolo, A., et al. High sensitivity mapping of brain-wide functional networks in awake mice using simultaneous multi-slice fUS imaging. Imaging neuroscience (Cambridge, Mass.) 1, 1–18 (2023).

53. Sauvage, J. et al. 4D Functional Imaging of the Rat Brain Using a Large Aperture Row-Column Array. IEEE Trans. Med. Imaging 39, 1884–1893 (2020).

54. Craig, A. D. How do you feel? Interoception: the sense of the physiological condition of the body. Nat. Rev. Neurosci. 3, 655–666 (2002).

55. Norman, S. L. et al. Single-trial decoding of movement intentions using functional ultrasound neuroimaging. Neuron 109, 1554–1566.e4 (2021).

56. Yoshiyama, M., Roppolo, J. R. & Groat, W. C. De. Alteration by urethane of glutamatergic control of micturition. Eur. J. Pharmacol. 264, 417–425 (1994).

57. Tiran, E. et al. Transcranial Functional Ultrasound Imaging in Freely Moving Awake Mice and Anesthetized Young Rats without Contrast Agent. Ultrasound Med. Biol. 43, 1679–1689 (2017).

58. El Hady, A., et al. Chronic brain functional ultrasound imaging in freely moving rodents performing cognitive tasks. J. Neurosci. Methods 403, (2024).

59. Dizeux, A. et al. Functional ultrasound imaging of the brain reveals propagation of task-related brain activity in behaving primates. Nature Communications 2019 10:1 10, 1400 (2019).

60. Demené, C. et al. Spatiotemporal Clutter Filtering of Ultrafast Ultrasound Data Highly Increases Doppler and fUltrasound Sensitivity. IEEE Trans. Med. Imaging 34, 2271–2285 (2015).

61. Maggi, C. A., Santicioli, P. & Meli, A. The nonstop transvesical cystometrogram in urethane-anesthetized rats: a simple procedure for quantitative studies on the various phases of urinary bladder voiding cycle. J. Pharmacol. Methods 15, 157–167 (1986).

62. Claron, J. et al. Large-scale functional ultrasound imaging of the spinal cord reveals in-depth spatiotemporal responses of spinal nociceptive circuits in both normal and inflammatory states. Pain 162, 1047–1059 (2021).

63. Foroosh, H., Zerubia, J. B. & Berthod, M. Extension of phase correlation to subpixel registration. IEEE Transactions on Image Processing 11, 188–199 (2002).

64. Ledoux, L. A. F., Brands, P. J. & Hoeks, A. P. G. Reduction of the clutter component in Doppler ultrasound signals based on singular value decomposition: a simulation study. Ultrason. Imaging 19, 1–18 (1997).

65. Pnevmatikakis, E. A. & Giovannucci, A. NoRMCorre: An online algorithm for piecewise rigid motion correction of calcium imaging data. J. Neurosci. Methods 291, 83–94 (2017).

66. Griggs, W. S. et al. Decoding motor plans using a closed-loop ultrasonic brain– machine interface. Nature Neuroscience 2023 27:1 27, 196–207 (2023).

67. Agyeman, K. A. et al. Functional ultrasound imaging of the human spinal cord. Neuron 112, 1710–1722.e3 (2024).

68. Chen, K., Kogan, J. F. & Fontanini, A. Spatially Distributed Representation of Taste Quality in the Gustatory Insular Cortex of Behaving Mice. Current Biology 31, 247–256.e4 (2021).

69. Friedrich, J., Giovannucci, A. & Pnevmatikakis, E. A. Online analysis of microendoscopic 1-photon calcium imaging data streams. PLoS Comput. Biol. 17, e1008565 (2021).

70. Stringer, C. & Pachitariu, M. Computational processing of neural recordings from calcium imaging data. Curr. Opin. Neurobiol. 55, 22–31 (2019).

71. Hickey, S. et al. Simultaneous T2 -weighted real-time MRI of two orthogonal slices. Magn. Reson. Med. 90, 2388–2399 (2023).

72. Jenkinson, M. & Smith, S. A global optimisation method for robust affine registration of brain images. Med. Image Anal. 5, 143–156 (2001).

73. Vercauteren, T., Pennec, X., Perchant, A. & Ayache, N. Diffeomorphic demons: efficient non-parametric image registration. Neuroimage 45, (2009).

74. Wolberg, G. & Zokai, S. Robust image registration using log-polar transform. IEEE International Conference on Image Processing 1, 493–496 (2000).

75. Zhong, S., et al. Comparative evaluation of image registration techniques in functional ultrasound imaging. Imaging neuroscience (Cambridge, Mass.) 3, (2025).

76. Goshtasby, A. Image registration by local approximation methods. Image Vis. Comput. 6, 255–261 (1988).

77. de Senneville, B. D., Manjon, J. V & Coupé, P. RegQCNET: Deep Quality Control for Image-to-template Brain MRI Affine Registration. Phys. Med. Biol. 65, (2020).

78. Cleveland, W. S. & Devlin, S. J. Locally Weighted Regression: An Approach to Regression Analysis by Local Fitting. J. Am. Stat. Assoc. 83, 596–610 (1988).

79. Czolbe, S., Pegios, P., Krause, O. & Feragen, A. Semantic similarity metrics for image registration. Med. Image Anal. 87, 102830 (2023).

80. Chang, C. & Glover, G. H. Time–frequency dynamics of resting-state brain connectivity measured with fMRI. Neuroimage 50, 81–98 (2010).

81. Sun, F. T., Miller, L. M. & D’Esposito, M. Measuring interregional functional connectivity using coherence and partial coherence analyses of fMRI data. Neuroimage 21, 647–658 (2004).

82. Welch, P. D. The Use of Fast Fourier Transform for the Estimation of Power Spectra: A Method Based on Time Averaging Over Short, Modified Periodograms. IEEE Transactions on Audio and Electroacoustics 15, 70–73 (1967).

83. Benjamini, Y. & Hochberg, Y. Controlling the False Discovery Rate: A Practical and Powerful Approach to Multiple Testing. Source: Journal of the Royal Statistical Society. Series B (Methodological) 57, 289–300 (1995).

84. Genovese, C. R., Lazar, N. A. & Nichols, T. Thresholding of Statistical Maps in Functional Neuroimaging Using the False Discovery Rate. Neuroimage 15, 870– 878 (2002).

85. Breiman, L. Random forests. Mach. Learn. 45, 5–32 (2001).

86. Mun, E.-Y. & Geng, F. Predicting post-experiment fatigue among healthy young adults: Random forest regression analysis. Psychol. Test Assess. Model. 61, 471 (2019).

87. Archer, K. J. & Kimes, R. V. Empirical characterization of random forest variable importance measures. Comput. Stat. Data Anal. 52, 2249–2260 (2008).

88. Strobl, C., Boulesteix, A. L., Kneib, T., Augustin, T. & Zeileis, A. Conditional variable importance for random forests. BMC Bioinformatics 2008 9:1 9, 307 (2008).

89. Fritsch, F. N. & Carlson, R. E. Monotone Piecewise Cubic Interpolation. SIAM J. Numer. Anal. 17, 238–246 (1980).

90. Breiman, L. Bagging Predictors. Machine-mediated learning 24, 123–140 (1996).

91. Liaw, A. & Wiener, M. Classification and Regression by randomForest. 2, (2002).

92. Freund, Y. & Schapire, R. E. Journal of Computer and System Sciences s SS1504 journal of computer and system sciences. 55, 1199139 (1997).

