## supplemental for "Distributed brain and spinal cord dynamics during micturition revealed by functional ultrasound imaging"

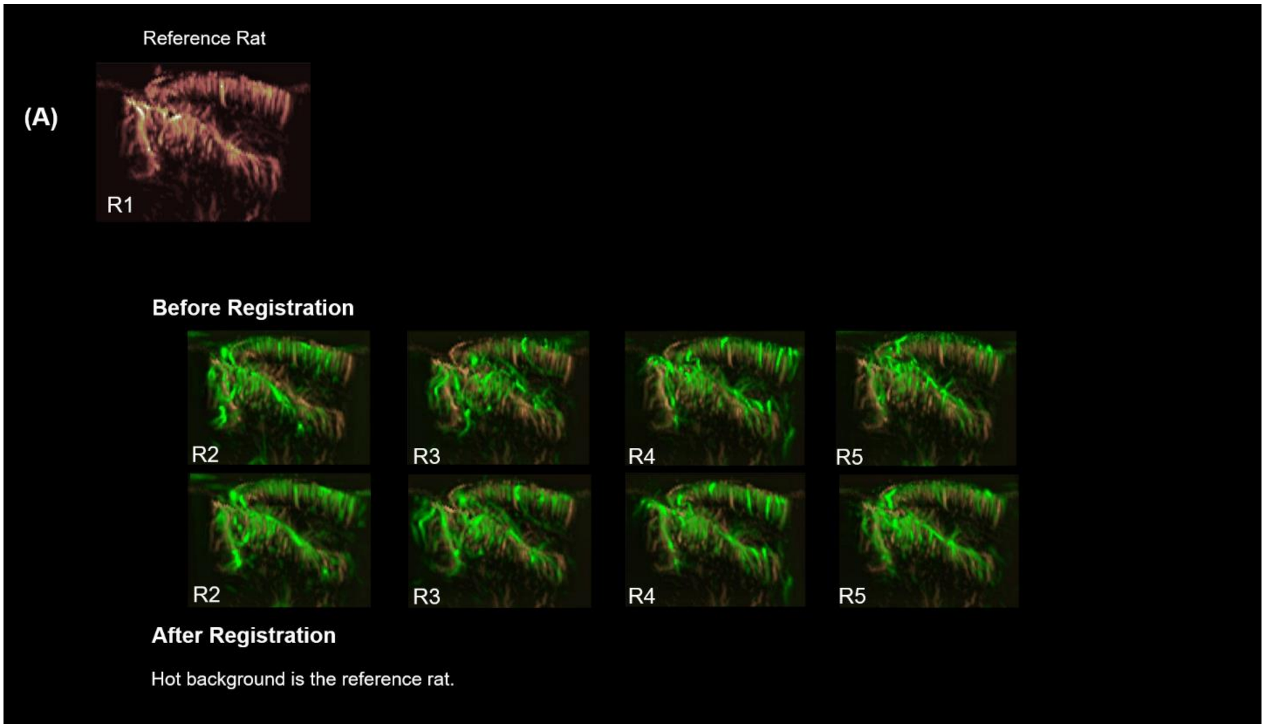


**Figure S1**. **Registration outcomes for brain pD images.** Individual brain pD vascular maps before and after affine registration to the reference rat (Rat 1). Pre- and post-registration images are overlaid onto the reference brain to evaluate alignment quality of vascular landmarks across animals.

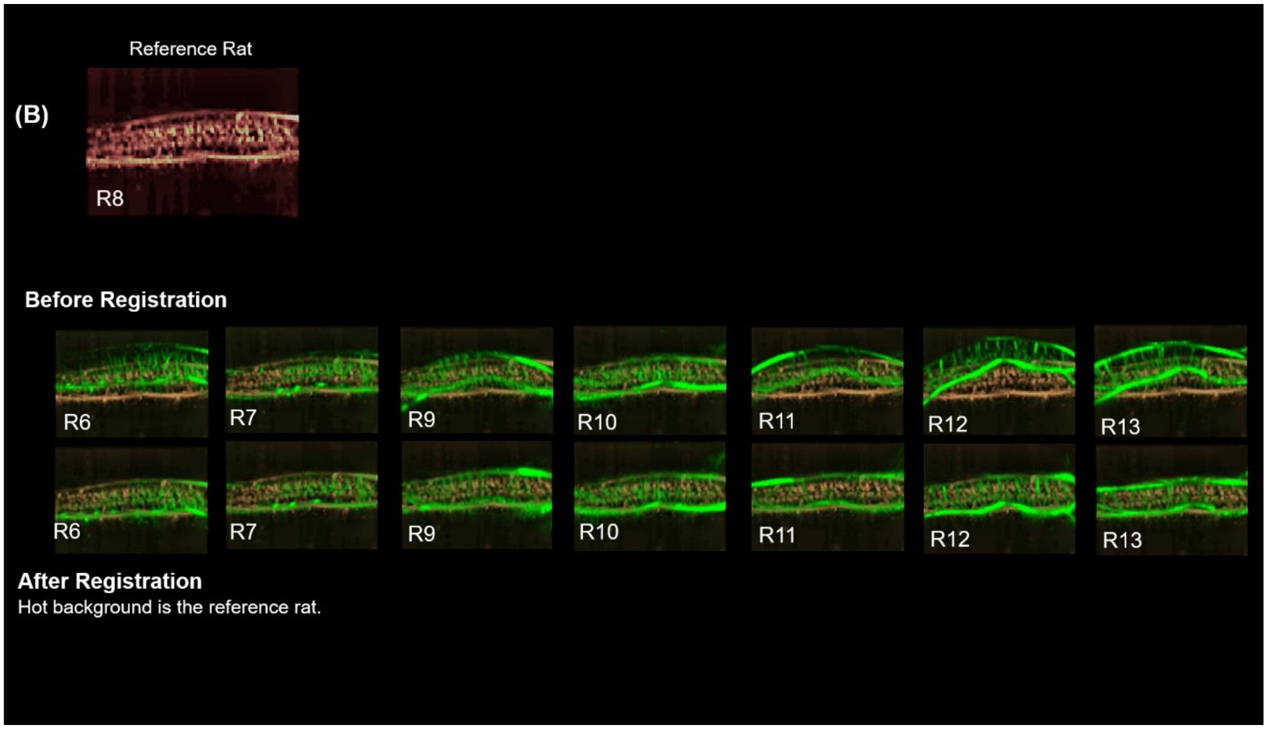


**Figure S2**. **Registration outcomes for spinal cord pD images.** Individual spinal cord pD maps before and after locally weighted mean (LWM) registration to the reference rat (Rat 8). Pre- and post-registration images are overlaid onto the reference spinal cord to evaluate cross-animal alignment quality.

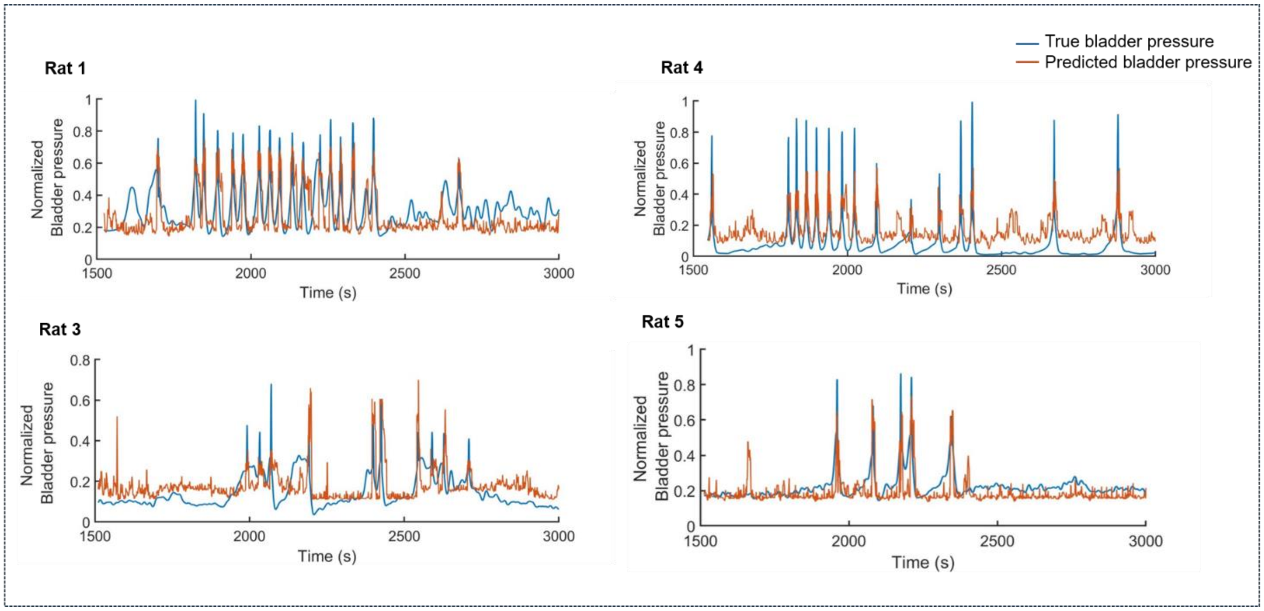


**Figure S3. Brain-group Random Forest Regression (RFR) predictions of bladder pressure for individual rats.** Predicted (orange) and true (blue) normalized bladder-pressure traces for the remaining four rats in the brain recording group. Predictions were generated using the final brain-model hyperparameters (100 learning cycles and a minimum leaf size of 50).

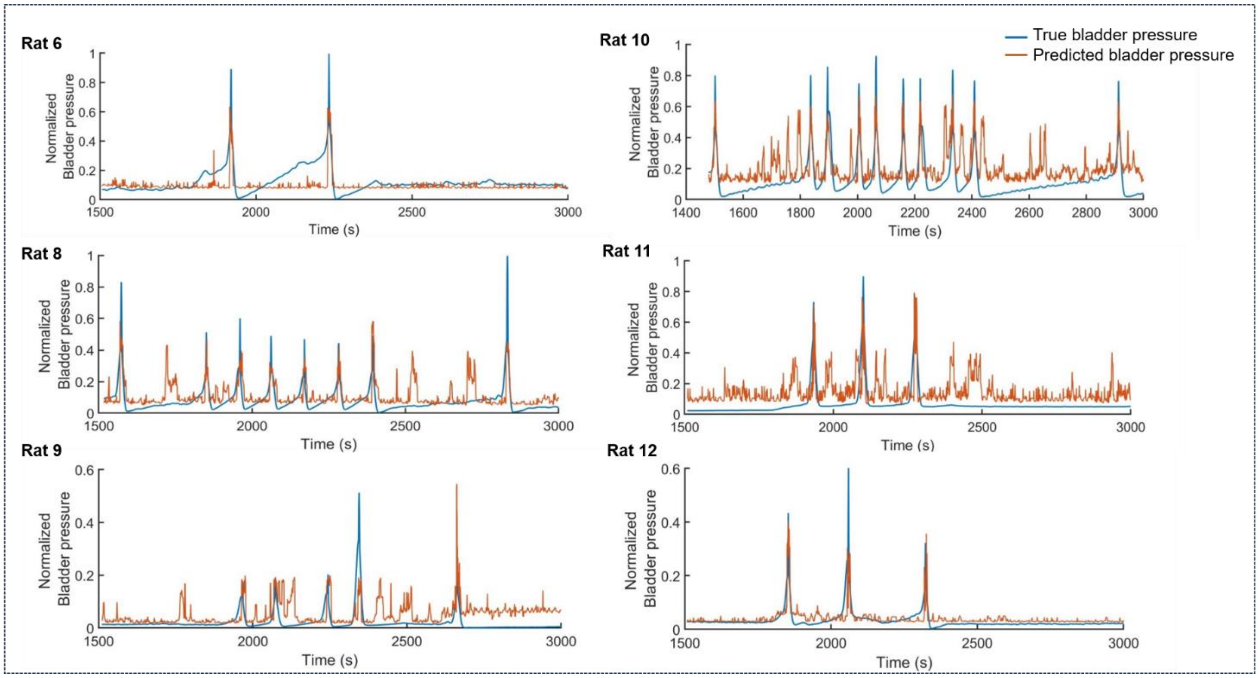


**Figure S4**. **Spinal cord group Random Forest Regression (RFR) predictions of bladder pressure for individual rats.** Predicted (orange) and true (blue) normalized bladder-pressure traces for the remaining six rats in the spinal cord recording group. Predictions were generated using the final spinal cord-model hyperparameters (500 learning cycles and a minimum leaf size of 100).

---

**Supplementary Movie 1. Spatiotemporal evolution of brain hemodynamics during the voiding cycle.**

Time-lapse of brain dynamic statistical parametric maps (dSPMs) illustrating cerebral blood volume changes (ΔCBV) around the voiding peak. dSPMs were generated using two-sided pixel-wise Student's t-tests ($p < 0.01$) with FDR correction, computed across overlapping 3-s sliding windows advancing in 1-s steps, relative to a 3-s reference period (−16 to −14 s relative to the voiding peak). For each pixel and time window, z-scores were calculated from percentage signal changes relative to this reference. Significantly positively (reddish regions) and negatively (bluish regions) modulated pixels are overlaid on the mean baseline pD intensity image of rat 3. Low-magnitude z-scores were rendered transparent to reduce visual crowding, and Gaussian smoothing ($\sigma = 1$) was applied for display.

---

**Supplementary Movie 2. Spatiotemporal evolution of spinal cord hemodynamics during the voiding cycle.**

Same as Supplementary Movie 1, but illustrating spinal cord hemodynamic changes ($\Delta$SCBV) around the voiding peak. dSPMs are overlaid on the mean baseline pD intensity image of the reference rat (rat 8).
